# Molecular basis of proteolytic cleavage regulation by the extracellular matrix receptor dystroglycan

**DOI:** 10.1101/2022.04.04.487063

**Authors:** MJM Anderson, AN Hayward, AT Smiley, K Shi, MR Pawlak, EJ Aird, E Grant, L Greenberg, H Aihara, RL Evans, C Ulens, WR Gordon

## Abstract

The dystrophin glycoprotein complex (DGC), anchored by the transmembrane protein dystroglycan, functions to mechanically link the extracellular matrix to the actin cytoskeleton to drive critical aspects of development and adult homeostasis. Breaking this connection via mutation of the actin adaptor protein dystrophin or impaired glycosylation of dystroglycan are strongly associated with diseases such as Muscular Dystrophy, yet cleavage of the dystroglycan protein by matrix metalloproteinases (MMPs) remains an understudied mechanism to disrupt the DGC. We solved X-ray structures of the membrane-adjacent domain of dystroglycan to understand the molecular underpinnings of dystroglycan MMP cleavage regulation. Dystroglycan proteolysis occurs within the versatile SEAL domain, which supports proteolysis in diverse receptors to facilitate mechanotransduction, protection of cell membranes, and even viral entry. The structure reveals a c-terminal extension of the SEAL domain that buries the MMP cleavage site by packing into a hydrophobic pocket, a unique mechanism of MMP cleavage regulation. We further demonstrate that structure-guided and disease-associated mutations disrupt proteolytic regulation using a new cell-surface proteolysis assay. Finally, we find that disruption of proteolysis leads to altered cellular mechanics and migration using high-throughput DNA tension probe and wound healing assays. These findings highlight that disrupted proteolysis is a relevant mechanism for “breaking” the DGC link to contribute to disease pathogenesis and may offer new therapeutic avenues for dystroglycanopathies.

## INTRODUCTION

Cells participate in bidirectional communication with their microenvironment to integrate biomechanical information and direct downstream responses such as cytoskeletal rearrangements and altered YAP localization (1, 2). In focal adhesions, transmembrane integrin receptors anchor a multi-protein complex that links the extracellular matrix (ECM) to the actin cytoskeleton to facilitate cell adhesion, contractility, and motility (3).

In parallel to integrins, the transmembrane receptor dystroglycan anchors the multi-protein dystrophin glycoprotein complex (DGC) to connect the ECM to the cellular cytoskeleton, most typically in cells that adjoin basement membranes, such as epithelial, neural, and muscle cells (4–8). Thus, dystroglycan plays important roles during development and adult homeostasis. For example, the critical mechanical link between the ECM and the actin cytoskeleton protects the sarcolemma from the forces of muscle contraction (9, 10), and dystroglycan expressed on astrocyte endfeet in the brain links to the basement membrane of endothelial cells lining blood vessels to help maintain the blood-brain barrier (11).

Disruption of the DGC mechanical link is strongly associated with disease. The connection between dystroglycan and actin is disrupted in Duchenne Muscular Dystrophy via mutations in dystrophin, the adaptor protein coupling dystroglycan to actin (12–14). Interestingly, analogous dystrophin mutations occur in 60-95% of myogenic tumors and are associated with worse outcomes, validating dystrophin as a tumor suppressor (15–17). Impaired glycosylation of dystroglycan has been shown to disrupt its linkage with laminin in the ECM, which has been observed in both muscular dystrophies and cancer (17–24). Mutations in dystroglycan itself are rare as its loss leads to embryonic lethality, but two mutations (T192M in the laminin-binding region and C669F in the membrane-adjacent SEAL domain) have been identified in patients with severe muscular dystrophy (8, 25–27).

However, there is an additional mechanism to break the DGC mechanical link that is less studied. Dystroglycan is known to be cleaved at a site just extracellular to the membrane by Matrix Metalloproteases such as MMP2 and 9 (11). In the brain, this occurs during axon guidance and myelination (28–31). There is some evidence that dystroglycan can be further processed by intramembrane cleavage, similar to Notch, to release intracellular fragments with distinct roles in the cell (32, 33). Given that MMPs are upregulated in contexts such as muscle disease, brain inflammation, and cancer, dystroglycan proteolytic fragments have been observed in many pathogenic contexts (34–38). Indeed, MMP inhibition decreased muscular dystrophy pathology and improved skeletal muscle function and regeneration in preclinical models (39–41). Thus, understanding how proteolysis is regulated in dystroglycan could lead to new therapeutic avenues.

MMP cleavage of dystroglycan occurs within a remarkably versatile membrane-adjacent domain called a SEAL (SEA-like) domain, named after the first three proteins it was found in sea-urchin Sperm protein, Enterokinase, and Agrin (42). This domain occurs in diverse mammalian receptors including Notch, ACE2, mucins, and protocadherins, and is characterized by a highly conserved core structure despite low sequence homology that features multiple sites of proteolytic cleavage. SEAL domains were bioinformatically identified in additional cell-surface proteins (43) and were also identified recently in bacteria as a domain that facilitates communication of mechanical stimuli (44). The domain facilitates proteolytic events important for mechano-signaling in Notch receptors (45, 46), epithelial tight junction barrier maintenance and repair via EpCAM (47), cellular membrane protection from shear force by mucins (48), and even viral entry by SARS-CoV2 via ACE2 (49–51).

Notch’s SEAL domain extensively interacts with its N-terminal cysteine-rich domain to compose the Negative Regulatory Region (NRR) that deeply buries a metalloproteinase cleavage site (45, 46). Forces associated with cell-to-cell contact induce a conformational change in the NRR to expose the protease site and trigger activation of the receptor (52). We recently demonstrated that membrane-adjacent SEAL domains from dystroglycan and several other receptors in cooperation with their N-terminal domains can functionally substitute for the Notch NRR in the context of synthetic Notch signaling assays, suggesting that dystroglycan proteolysis might be regulated by a similar mechanism as Notch receptors by restraining proteolysis in the absence of mechanical forces (53).

Thus, we set out to understand the molecular underpinnings of dystroglycan proteolytic regulation and the potential for proteolysis to disrupt cell mechanics mediated by the DGC mechanical link. The X-ray structure of the dystroglycan proteolysis domain comprised of tandem immunoglobulin-like (IGL) and SEAL domains shows that the MMP site is occluded in the structure, but by a structural mechanism distinct from Notch receptors and other SEAL domain-containing receptors. Moreover, we show that disease-associated and structure-guided mutations lead to increased protease sensitivity. Finally, we also demonstrate that proteolysis decreases cellular traction force in a similar manner to muscular dystrophy-associated dystrophin mutations.

## RESULTS

### Characterization of dystroglycan “proteolysis domain” in secreted and full-length receptors

Prior to structural studies, we characterized the tandem IGL and SEAL domains to further understand the requirements for correct domain folding and susceptibility to MMP cleavage. For domain folding, we assessed the propensity of the constructs to undergo correct autoproteolysis during expression. A feature of the SEAL domains found in mucins and dystroglycan is an autocatalytic processing cleavage that occurs during trafficking which results in non-covalently associated subunits at the cell surface (48, 54). Previous studies into this autoprocessing of mucins have demonstrated that cleavage requires proper domain folding, which positions and prepares a nucleophilic serine for self-proteolysis in a highly constrained loop of the SEAL domain (48). Autoproteolysis thus results in two separate bands on an SDS-PAGE gel, providing a convenient readout of correct folding. We secreted dual epitope-tagged dystroglycan-Fc fusion constructs featuring various truncations and mutations, purified them on Protein A beads, subjected them to MMP-mediated cleavage, and performed immunoblots to determine the extent of autoproteolysis, which is reflective of proper domain folding (Figure 1A). In comparing the processing of dystroglycan truncations, we found that constructs containing both the IGL and SEAL domains were completely processed, whereas truncations containing the SEAL domain alone had a substantial reduction in this processing (Figure 1B), implying the IGL domain aids in proper folding/trafficking of the SEAL domain. Treatment of the constructs with exogenous MMPs (either MMP-2 or MMP-9) demonstrates that the intact IGL+SEAL domain is resistant to MMP treatment while the isolated SEAL domain is susceptible to cleavage. This suggests MMP cleavage is regulated by the conformation of the domain and shows that the SEAL domain lacking its adjacent IGL domain is defective in either establishing or maintaining the proper tertiary structure required for MMP cleavage regulation. This reduced autoprocessing and increased susceptibility to MMPs is also observed in the context of the muscular dystrophy-associated point mutations, I593D and C669F (Figure 1C, Figure S1A) (26, 55), which suggests that these mutations may mediate their disease phenotype by diminishing proper domain folding of dystroglycan. The muscular dystrophy-associated mutation, S654A, eliminates the catalytic hydroxyl needed for autoprocessing yet does not increase susceptibility to MMP cleavage (56). Previous research shows that the corresponding mutation in other SEALs destabilizes some protein structures, but not all (discussed below). How S654A causes muscular dystrophy in mice is still uncertain, though it points to an important, undiscovered role of autoprocessing. Finally, mutations and truncations within the IGL and SEAL domains have similar effects on both autoprocessing and MMP cleavage susceptibility in the context of full-length dystroglycan (Figure S1B). Together, these data show the IGL and SEAL domains cooperate to form a functional “proteolysis domain” that supports the correct folding necessary for autoprocessing and resisting MMP cleavage.

**Figure 1.**
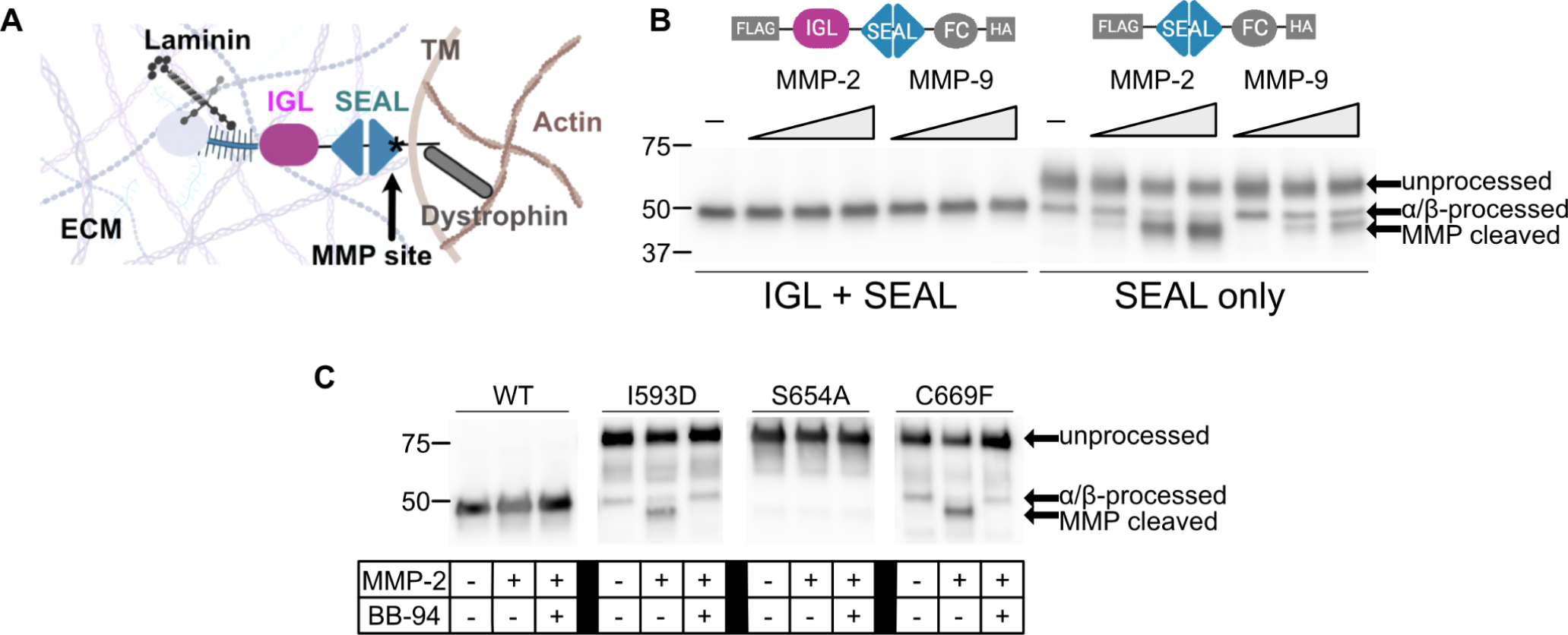
Intact dystroglycan proteolysis domains resist MMP cleavage but truncated or mutated domains become sensitive to MMPs. **A)** Schematic of dystrophin-glycoprotein complex (DGC) and dystroglycan domains. **B)** Anti-HA Western blot of dystroglycan expression constructs exposed to increasing amounts of MMP-2 and MMP-9. Concentrations of MMP-2 and MMP-9 are 0.07 ug/mL, 0.37 ug/mL, and 0.73 ug/mL. **C)** Anti-HA Western blot of dystroglycan expression constructs with muscular dystrophy associated mutations, exposed to 0.73 ug/mL +/− 40 uM BB-94 (Batimastat, MMP inhibitor).

### Molecular basis of proteolytic regulation

We next used X-ray crystallography to solve a 2.4 Å structure of the dystroglycan tandem IGL and SEAL domains to determine the molecular basis of the conformational regulation of dystroglycan proteolysis (Table 1). The asymmetric unit is composed of two copies of the dystroglycan proteolysis domain, with density for residues G491-V722 in one structure and E492-P720 in the other (Figure 2A). The two molecules in the asymmetric unit interact in an antiparallel fashion and align with 0.528 Å RMSD when overlaid. The overall monomer structure shows the IGL and SEAL domains interacting to form an L-shape. The N-terminal domain IGL domain shows a characteristic two-layer Ig-domain β-sandwich and the C-terminal SEAL domain is characterized by a **βαββαβ** sandwich as observed in other SEAL domain structures (Figure 2B). Both molecules bind to a Ca^+2^ ion near the N-terminus and a Cl^−^ between the IGL and SEAL doamains. This structure has been deposited in the Protein Data Bank (PDB) as 8UF4.

**Figure 2.**
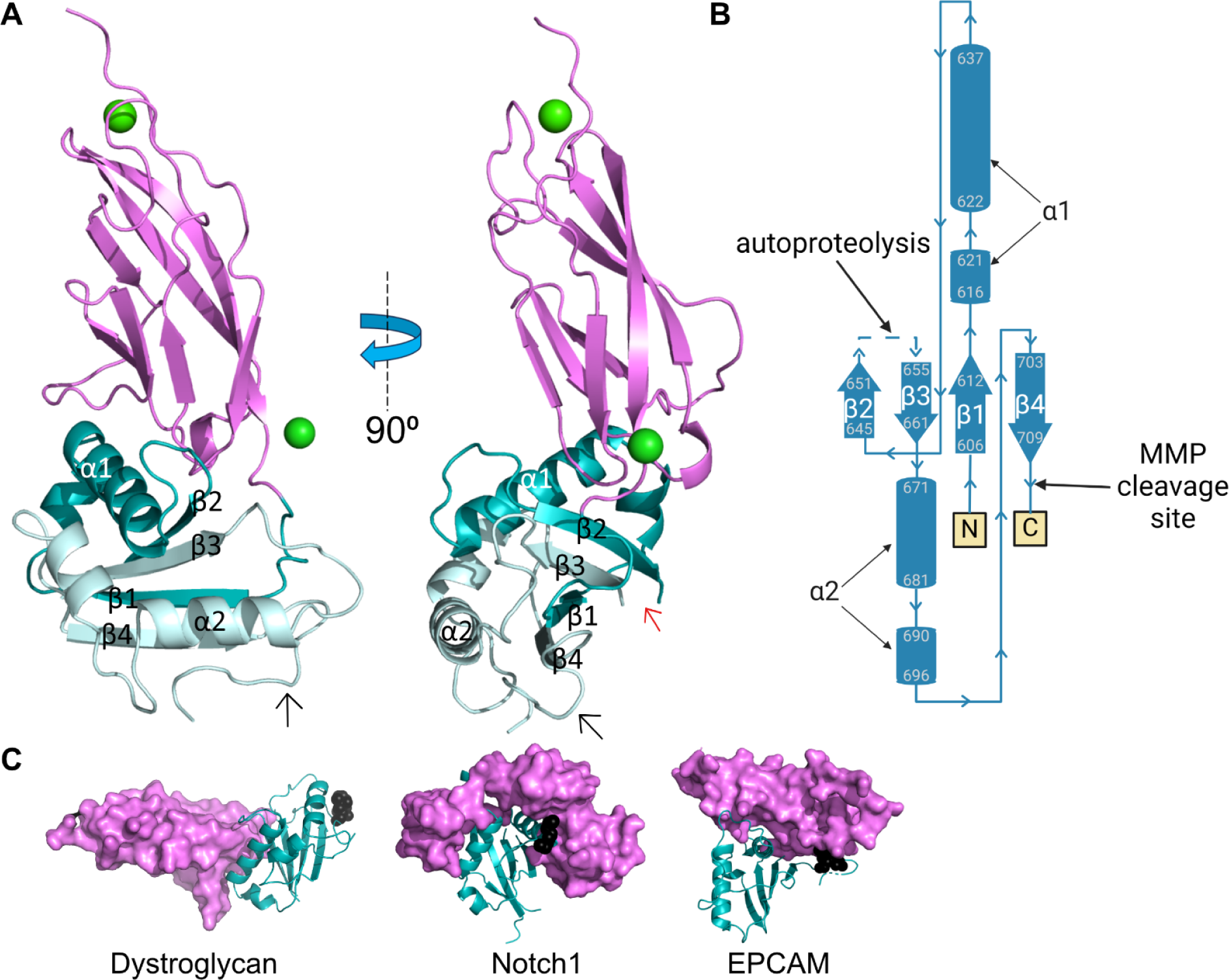
Dystroglycan X-ray crystallography structure reveals key differences from other SEAL domains. **A)** Cartoon representation of dystroglycan IGL (magenta) and SEAL N-terminal to autoproteolysis site (cyan) and autoproteolysis site to C terminus (greencyan). The autoproteolysis site (red) and MMP cleavage site (black) are pointed out with arrows. **B)** Topology map of dystroglycan SEAL. **C)** Surface representation of SEAL (cyan), MMP cleavage site (black), and neighboring domains (magenta) from dystroglycan, Notch1 (PDB: 3I08) and EPCAM (PDB: 4MZV).

The dystroglycan structure shows some interesting similarities and differences when compared to other SEAL domain-containing structures (Table 2, Table S1). The overall SEAL **βαββαβ** architecture shows a high degree of structural conservation among diverse SEAL domains despite low sequence similarities. The SEAL domain of dystroglycan has RMSDs ranging from 2 to 7.5 Å with other known SEAL domain structures (1-7.5 if expanded to predicted structures of SEALS). The high structural homology of SEAL domains is also notable given that the domain supports different mechanisms of receptor processing: dystroglycan and mucins are autoprocessed while the Notch and Agrin SEAL domains are proteolyzed in the same location in a large unstructured loop by exogenous proteases (48, 54, 57, 58). Other SEAL domains are not known to be processed into noncovalently linked subunits.

The most obvious difference among SEAL domains is their relative interaction with adjacent N-terminal domains (Fig 2C, SI Fig 2). Several SEAL domains do not contain structured N-terminal domains, such as mucins. Other N-terminal domains are structurally diverse (Table 2, Table S1). SEAL domains from Notch and EpCAM interact extensively with adjacent domains as quantified by having over 1500 Å^2^ of buried surface between the SEAL and adjacent domains. These interactions are important for protein function as the adjacent domain covers an ADAM-family metalloprotease cleavage site which creates the structural basis for proteolytic cleavage regulation in Notch, as the adjacent must be disengaged by mechanical forces for cleavage to occur (45, 46, 52). The SEAL domains from dystroglycan and others interact less extensively (400-800 Å^2^ buried surface area) with interactions forming far from the MMP cleavage site. Thus, dystroglycan regulates MMP cleavage in a distinct mechanism from previously studied SEAL domains.

We hypothesized that dystroglycan may form a dimer as part of its natural function because of high structural homology between its proteolysis domain and the membrane-associated domain (MAD) of Protocadherin-15 (PCDH15) (Figure S3A), which was recently shown to form an obligate homodimer for proper protein function (59–63). We performed analytical ultracentrifugation (AUC) of expressed and purified dystroglycan proteolysis domain to investigate this and determined that dystroglycan does not form a dimer in solution (Figure S3B). Though our refined X-ray crystallography structure does contain a crystallographic dimer, it has a considerably smaller dimeric interface in comparison to PCDH15 which hints at a distinct difference in function despite high structural similarity.

One unique aspect of the dystroglycan structure in comparison to all of the others referenced thus far is a C-terminal SEAL domain extension which nestles in a hydrophobic pocket formed by **α**2 and **β**4 at the C-terminus of the SEAL domain (Figure 3). In all other SEAL domains, the C-terminus is unstructured, extending from the SEAL domain without making any contact (Figure S4). The amino acids of this extension (FIP718-720) include the mapped MMP cleavage site (HL715-716) and a disulfide bond formed between Cys669 and Cys713. Notably, Cys669 is mutated in a patient with severe muscular dystrophy, indicating that disruption of this bond results in dysregulation of proper proteolysis domain folding (26, 64). We hypothesized that the disulfide bond and hydrophobic residues together act to anchor the MMP cleavage site into a hydrophobic pocket in the protein and inhibit cleavage. To our knowledge, no other SEAL domain is known to regulate MMP proteolysis through this mechanism, highlighting the importance of continuing to study this unique domain.

**Figure 3.**
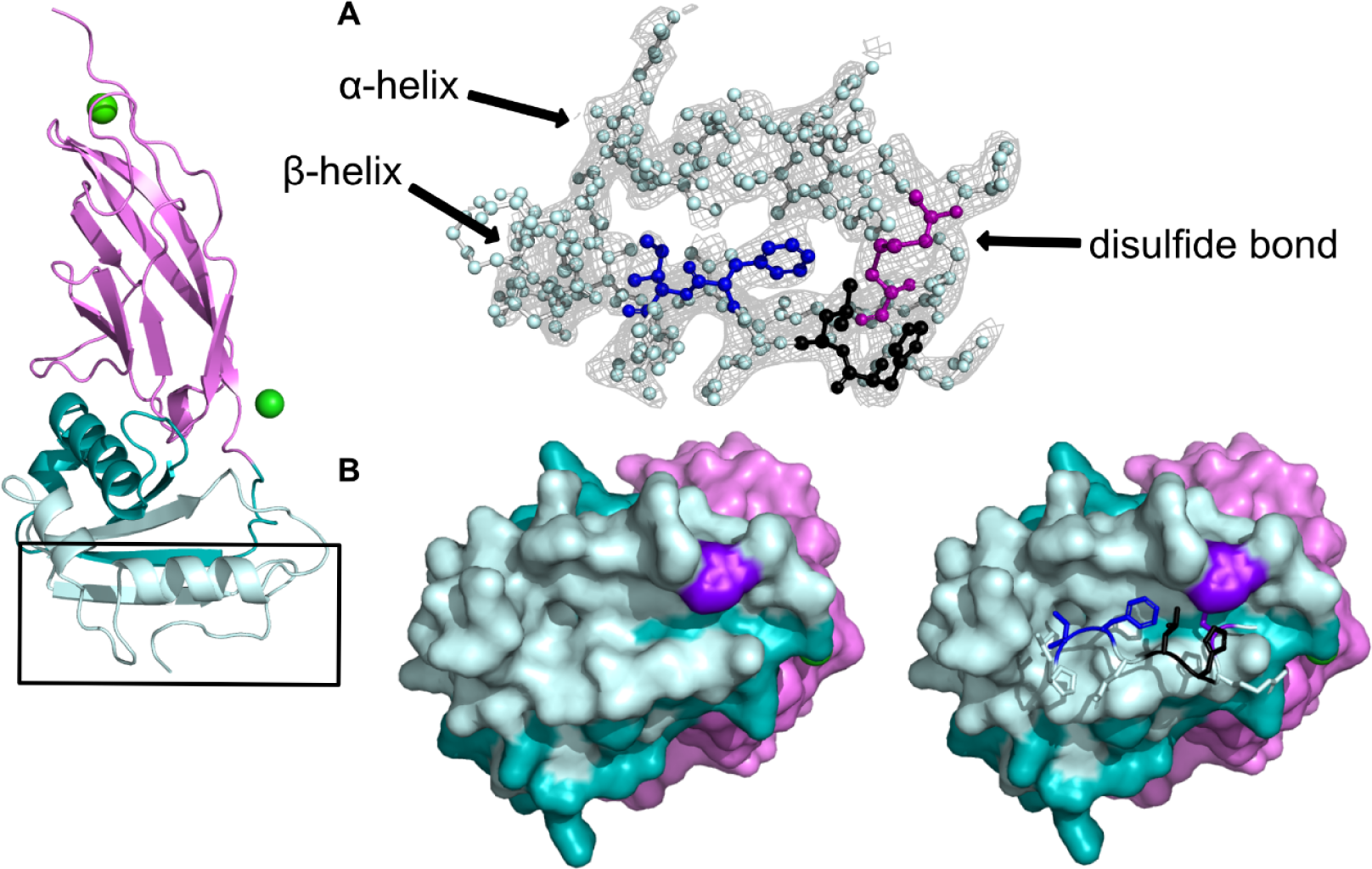
Dystroglycan C-terminus stabilizes MMP cleavage site. **A)** Electron density map with the refined structure, highlighting disulfide between Cys669 and Cys713 (purple), FI718-719 (blue), and MMP cleavage site (black). **B)** Hydrophobic pocket shown with (right) and without (left) SEAL C-terminal extension.

To understand the importance of these amino acids in the function of dystroglycan, we explored the evolutionary conservation of both the SEAL C-terminus and autoproteolysis site in dystroglycan. Previously, Adams and Brancaccio (65) identified representative dystroglycan homologs from all the phyla in which it is found. We used these sequences to calculate simple conservation scores and create logo plots to determine the evolutionary conservation of these regions of dystroglycan (Figure S5 top). We also calculated the conservation of these regions for the more closely related Chordata phylum which includes lancelets, sea squirts (and related species), and all vertebrates (Figure S5 bottom). Overall, the Chordata phylum shows a high level of conservation throughout the entire protein with the autoprocessing motif, GSIVV, almost completely conserved, again highlighting the importance of autoprocessing for proper domain folding. The notable FIP718-720 C-terminal extension is almost completely conserved in the Chordata phylum, with Valine occasionally found in place of Ile719, suggesting that having hydrophobic residues in this part of the protein are important for function. These trends are similar though, not as well conserved, when looking at the entire evolutionary time of dystroglycan, with GSIVV being the dominant sequence at the autoproteolysis site and FIP718-720 (or other combinations of comparable hydrophobic residues) being conserved in an area with generally low conservation. Surprisingly, Cys713 and Cys669 (not shown) are completely conserved throughout all species, highlighting the importance of this disulfide bond for proper function of the C-terminal extension, and perhaps for stability of the proteolysis domain overall. This consistently high evolutionary conservation, paired with our structural data, indicates that these regions are functionally important, with putative roles in the maintenance of tertiary structure that regulates MMP cleavage.

### Structural disruption of C-terminal extension packing increases cell surface proteolysis

To test our hypothesis that dystroglycan MMP cleavage is regulated by anchoring of of the C-terminal extension against the SEAL domain, we made multiple mutations of the structurally-implicated F718 (to A, R, and H) and C713 (to A and S). We expect these F718 mutations to disrupt the hydrophobic interactions normally made by this amino acid, thus exposing the MMP cleavage site and increasing MMP susceptibility. Similarly, the cysteine mutations will break the disulfide bond (like the muscular dystrophy mutation, C669F), but without dramatically changing residue size or charge. We first tested the effects of these structure-guided mutations on cell surface proteolysis of dystroglycan in U251 cells due to their robust expression of MMPs and dystroglycan’s important role in glioblastoma pathology (66, 67). Because immunoblots are unable to differentiate between protein on the cell surface and protein that has been improperly trafficked (as some dystroglycan mutations are known to do), we looked to create an assay that would only readout cell-surface dystroglycan MMP cleavage in high throughput. To do this, we inserted Flag and HA epitope tags on either side of the proteolysis domain and transfected these constructs into U251 cells (Figure 4A). In the event of MMP cleavage, the Flag tag would be cleaved off with the majority of the dystroglycan ectodomain, decreasing binding by fluorescently-labeled anti-Flag antibodies, and therefore, fluorescence as readout by flow cytometry (Figure S6A left). However, the HA epitope tag is found on the small part of the dystroglycan ectodomain that is retained even after MMP cleavage, allowing it to act as a marker of overall cell surface dystroglycan expression by binding to fluorescently labeled anti-HA antibodies (Figure S6A right). A representative contour plot of both values shows the shift in APC fluorescence without a large change in FITC fluorescence (Figure 4B). Since cells do not need to be permeabilized for this assay, it only assesses the cleavage of dystroglycan found on the cell surface.

**Figure 4.**
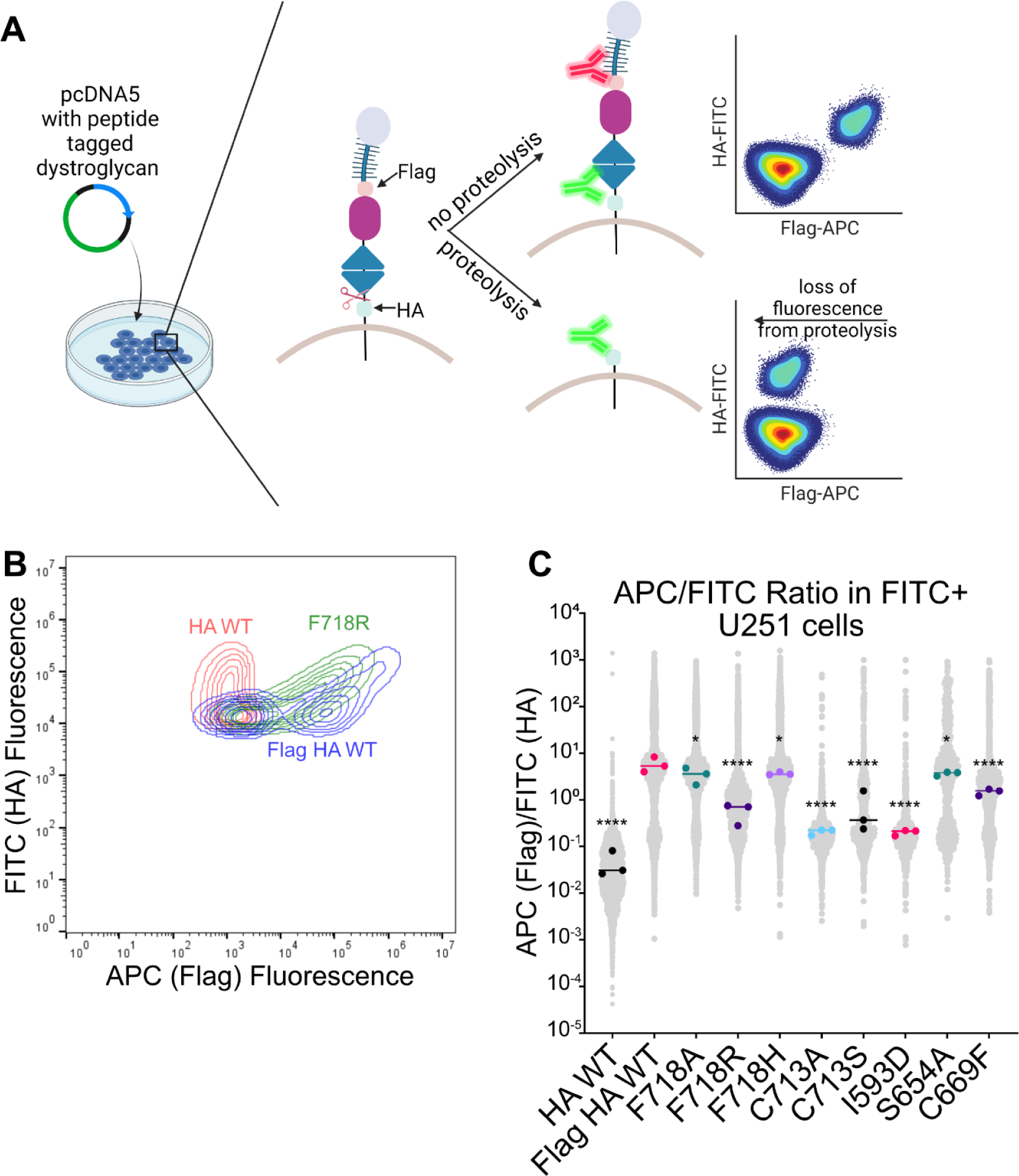
Dystroglycan mutation increases cleavage by MMPs. **A)** Schematic representation of experiment. **B)** Contour plot of representative samples to show where loss of flag tag causes shift in APC fluorescent value. HA WT is a control which does not have a Flag tag present. **C)** APC/FITC ratio for structure-guided and muscular dystrophy-associated mutations in FITC+ U251 cells. Large dots represent median for individual days and the line represents overall median. Significance was calculated using a one-way ANOVA with follow-up Dunnett’s multiple comparisons test to Flag HA WT. ns = nonsignificant, * = p<0.05, ** = p<0.01, *** = p<0.001, **** = p<0.0001.

To analyze these data, we used the ratio of APC (Flag tag, lost if cleaved)/FITC (HA tag, expression control) fluorescent signal to compare MMP proteolysis of structure-guided and muscular dystrophy mutations of dystroglycan (Figure 4C). All tested mutations show a decreased ratio compared to wildtype, which suggested increased MMP cleavage. A similar pattern was seen in other cell lines (NIH-3T3, COS-7, HAP-1 DAG1KO, HEK293T) showing that these mutations affect protein cleavage regardless of cell type (Figure S6B). Surprisingly, S654A caused a mild increase in proteolysis, in contrast to what was seen in the immunoblots. This difference may come from increased sensitivity or from only analyzing cell surface dystroglycan. Regardless, disrupting the packing of the C-terminal extension via mutation exposes the MMP cleavage site and increases susceptibility to MMP cleavage at the cell surface, supporting the hypothesis of the unique shielding mechanism for MMP cleavage regulation found in dystroglycan.

### Modulating cell surface proteolysis correlates with altered cell mechanics

We next wanted to test the effects increased MMP cleavage has on dystroglycan’s mechanical function as a link between the ECM and the actin cytoskeleton. We hypothesized that alterations in MMP cleavage, which act to break adhesive contacts, will reduce the forces that cells exert on their surroundings. Indeed, we recently developed and applied a genetically-encoded vinculin tension sensor to demonstrate that dystrophin mutations reduced vinculin tensions and cellular traction forces in C2C12 myoblasts via cross talk between DGC and integrin adhesive complexes (2). Genetically-encoded tension sensors have wide application in the characterization of mutation-induced changes in cellular force and tension, or “mechanotype”, which is emerging as an important player in disease progression. However, these methods are often low throughput and labor intensive to image and analyze cells. Therefore, we recently developed a widely applicable “off-the-shelf” tension sensor system called Rupture and Deliver Tension Gauge Tethers (RAD-TGTs), which provide a straightforward means to characterize the mechanotype of cells in high throughput without need for cellular imaging (68). Briefly (Fig 5A), this method involves immobilizing well-characterized TGT DNA duplex(72) tension probes to standard 96-well cell culture plate via biotin-streptavidin chemistry. The TGTs encode a Cy5 fluorophore and are covalently linked to the high-affinity integrin ligand echistatin via a DNA-linking HUH-fusion tag(69–71) called WDV. The rupture force of the DNA duplexes is tunable between 12 and 54 pN (72). Cells are then plated on these probes, and will “rupture and deliver” into the cell a fraction of the DNA duplexes depending on cellular forces associated with adhesion, contractility, and motility. Thus flow cytometry can be used as a relative readout of cellular mechanotype, where cells exerting higher forces on its surroundings will rupture more RAD-TGTs and exhibit higher fluorescence than cells exerting lower forces.

**Figure 5.**
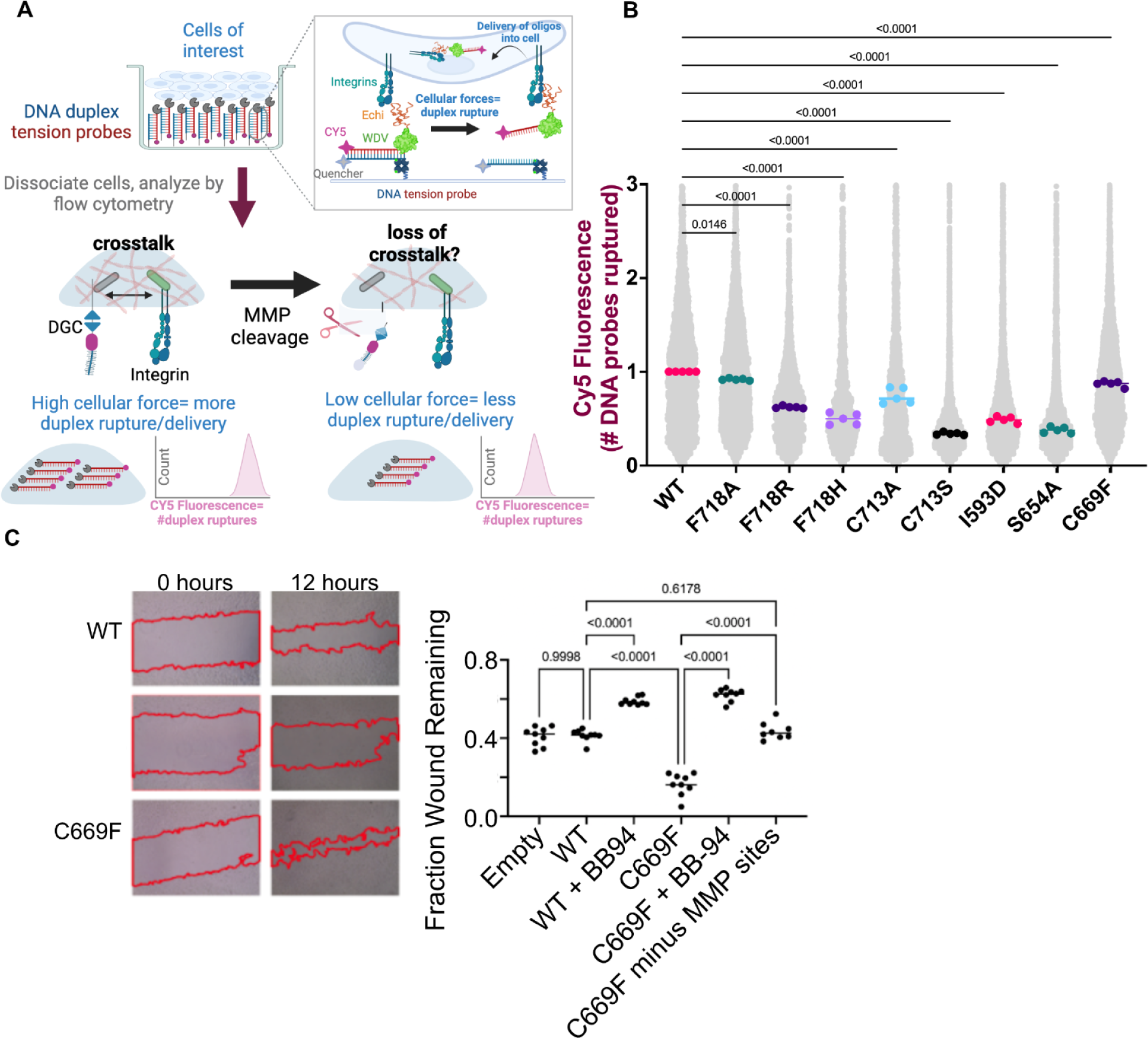
Dystroglycan mutation alters cellular mechanics and migration. **A)** Schematic representation of RAD TGT experiment. Echi-high-affinity integrin ligand echistatin; WDV-HUH-fusion tag linking ligand to DNA probe; DGC-dystrophin glycoprotein complex; MMP matrix metalloprotease; bb-94 batimistat protease inhibitor **B)** Relative Cy5 fluorescence compared to the median of WT for that day for rupture of 54pN RAG-TGTs. Large dots represent median for biological replicate and line represents overall median. Grey dots show the fluorescence of each cell across all biological replicates. Significance was calculated via one way ANOVA with follow-up Dunnett’s multiple comparisons test compared to WT. **C)** Representative images (left) of wound at 0 and 12 hours with quantification of wound remaining (right). Dots represent individual wells with the line representing overall median. Significance was calculated using t test.

To assess how modulating MMP-mediated cleavage of dystroglycan impacts cellular mechanics (i.e. mechanotype), we transiently transfected cells with dystroglycan variants with a C-terminal GFP fusion partner. Twenty-four hours post transfection, cells were plated onto CY5 labeled TGTs presenting the strong integrin ligand echistatin tethered to a fibronectin-coated glass plate for 2-4 hours after which cells are dissociated from the plate and analyzed via flow cytometry. In U251 cells, there is a statistically significant decrease in cellular forces in the context of structure guided and muscular dystrophy-associated mutations for both low and high molecular forces (Figure 5B, S7A). This trend holds in other cells as well, though not all were statistically significant (Figure S7), perhaps due to variation in the expression of cell surface integrins. Furthermore, gating solely on GFP+ cells does not affect the overall trend of decreased cellular mechanics (Figure S8). Therefore, dystroglycan MMP cleavage through mutations affects the mechanotype of cells, reducing the amount of force it exerts on the surroundings. This is consistent with prior observations on effects of muscular dystrophy associated mutations encoded in dystrophin, which links dystroglycan to the actin cytoskeleton. Those studies showed that mutations decreased vinculin tension and cellular traction force. These effects on cellular mechanics are likely due to crosstalk via the actin cytoskeleton, where the presence of an intact DGC enhances the actin cytoskeleton mesh and rendering the cells more contractile. Mutations that result in cleavage of dystroglycan disrupt the crosstalk and reduce cellular forces.

We next asked if MMP cleavage-sensitizing mutants altered cell migration. We chose to focus on the C669F disease-associated mutation and made an additional construct where we deleted the C-terminal extension of the SEAL domains which we reasoned would counteract the effects of the excess MMP cleavage caused by the C669F mutation. We also treated cells with the broad-spectrum metalloprotease inhibitor, BB-94, to generally block MMP proteolysis. We performed scratch assays in transiently transfected 3T3 cells and measured the propensity of cells to close wounds at different time points (Figure 5C). Blocking proteolysis via BB-94 treatment clearly slows down wound closure. In contrast, cells expressing MMP cleavage-sensitizing C669F mutation exhibit significantly faster rates of wound closure, an effect that is reversed when those cells are treated with BB-94. Finally, cells transfected with the C669F mutation in concert with removal of the MMP sites close wounds much closer to wildtype rates, as we predicted.

### Cancer-associated mutations similarly disrupt proteolytic regulation and cell mechanics

Having now shown that both muscular dystrophy and structure-guided mutations disrupt dystroglycan proteolytic regulation and cause altered cell mechanics, we wondered if a similar phenotype would be found with cancer mutations, as the DGC is commonly altered in a variety of cancers. We searched the Catalog of Somatic Mutations In Cancer (COSMIC) for mutations in the dystroglycan proteolytic domain and chose a subset to test (Figure 6A). Immunoblotting showed that most of the tested mutations increased dystroglycan proteolytic cleavage, similar to the muscular dystrophy and structure-guided mutations (Figure 6B, S9). Three of these mutations were then tested using flow cytometry for both proteolysis and cellular mechanics as above. While all three mutations showed a significant decrease in cellular force, only one had a statistically significant increase in proteolysis on flow cytometry in U251s (Fig 6C-D, S10-12). Thus, while cancer-associated mutations have a similar change in cellular mechanics as in cells with muscular dystrophy mutations, proteolytic cleavage seems to be unchanged. Further investigation is needed to elucidate the mechanism of cellular force aberration in cells with these mutations.

**Figure 6.**
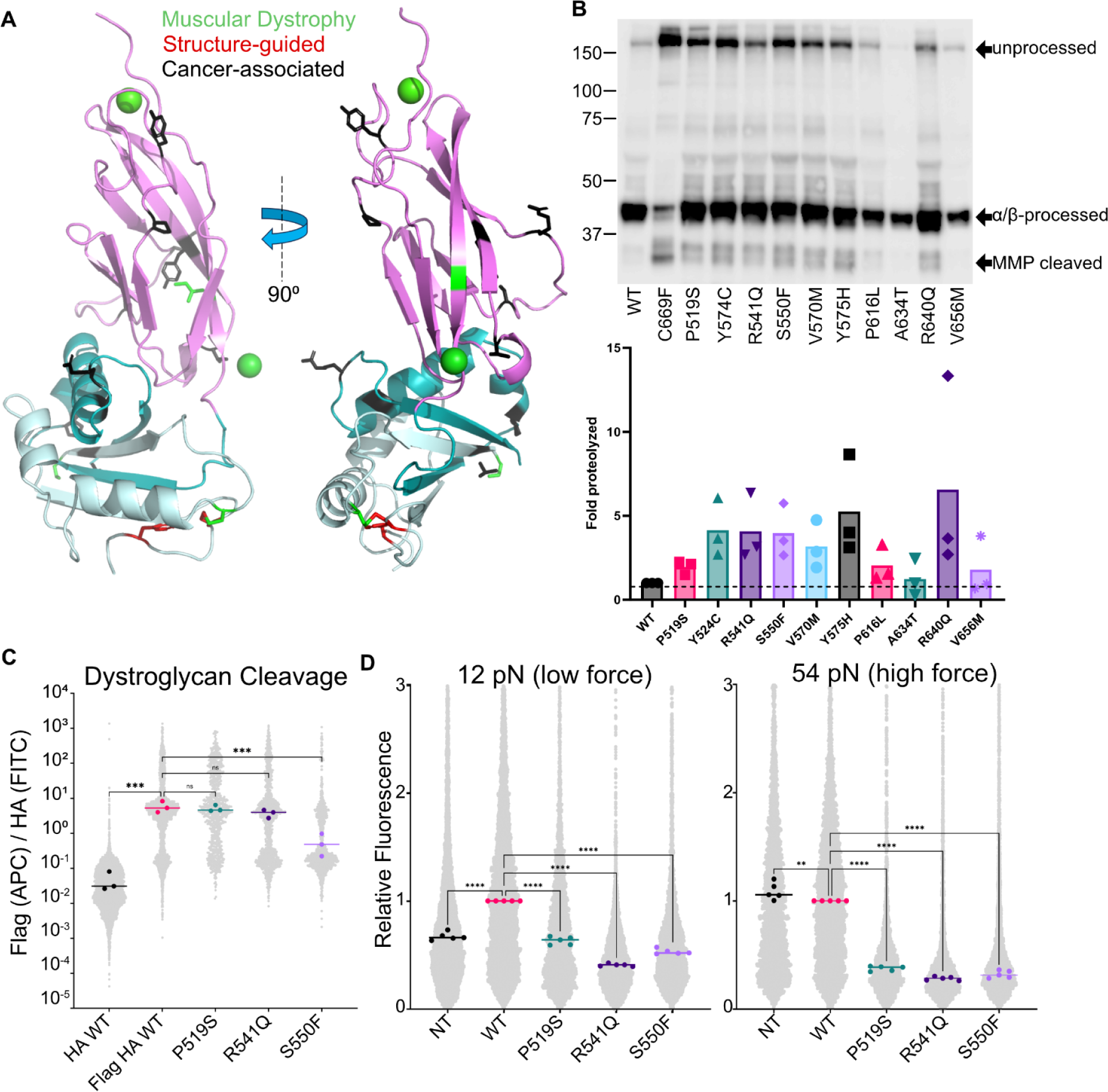
Cancer-associated mutations of dystroglycan alter MMP cleavage and cell mechanics. **A)** Cartoon representation of dystroglycan with muscular dystrophy (green), structure-guided (red), and cancer-associated (black) mutations colored. **B)** Anti-β-dystroglycan Western blot (top) with fold proteolysis compared to WT (bottom). Symbols represent individual blots with bar representing mean. **C)** APC [Flag]/FITC[HA] ratio for cancer-associated mutations in FITC+ U251 cells. **D)** Relative Cy5 fluorescence compared to the median of WT for that day as readout by RAD TGTs. (C-D) Large dots represent median for individual days and line represents overall median. Significance was calculated using a one-way ANOVA with followup Dunnett’s multiple comparisons test to Flag HA WT. ns = nonsignificant, * = p<0.05, ** = p<0.01, *** = p<0.001, **** = p<0.0001.

## DISCUSSION

The structurally conserved SEAL domain provides a hub for proteolytic cleavages and interactions with adjacent domains that enable diverse biological functions such as signaling, cellular plasticity, viral entry, and communication of mechanical stresses. In this study, we investigated the molecular basis of proteolytic cleavage regulation in the extracellular matrix receptor dystroglycan and how disease-associated and structure-guided mutations dysregulate proteolysis and affect cellular mechanics.

One proteolytic event commonly observed in SEAL domains occurs during receptor trafficking and results in non-covalently associated subunits at the cell surface. This proteolysis can be autocatalytic, like in mucins and bacterial SEAL domains or cleaved by exogenous proteases like in Notch. Recombinantly expressed dystroglycan is indeed split into two subunits without the addition of exogenous proteases and contains the canonical autoproteolytic site. Moreover, the IGL domain is required to stabilize the SEAL domain to facilitate this autocleavage, in contrast to mucins which do not have a structured adjacent domain.

The other type of proteolysis commonly occurring in SEAL domains has been mapped to multiple locations within the domain and occurs through metalloproteases. For example Notch, EpCAM, and ACE2 have cleavage that map to the C-terminus of the SEAL domain. It is still unclear how recognition of this site is determined. MMPs are known to be promiscuous in the amino acid sequences it will cleave while ADAMs (a disintegrin and metalloproteinase), including ADAM10 which cleaves Notch, cleave a certain distance from the membrane (73, 74). Regulation of this cleavage, therefore, is generally encoded by the SEAL-containing protein. In Notch the cleavage is tightly regulated by the adjacent domain, which covers the ADAM cleavage site. But in dystroglycan, the adjacent domain is on the opposite side of the SEAL domain and unable to regulate cleavage. Intriguingly a C-terminal extension off the SEAL domain that is highly conserved evolutionarily, pins the MMP cleavage site against the protein. This has not been observed in other SEAL domains where the C-terminus has no contact with the SEAL domain. Mutations that disrupt this packing lead to increases in MMP cleavage, and increases in cleavage lead to decreases in cellular force and increases in cell migration in wound healing assays.

The role of autoproteolysis and MMP cleavage are not well understood in dystroglycan. After autoproteolysis, the resulting subunits are associated by tightly interwoven beta strands in the SEAL domain. In mucins, dissociation of the two non-covalently associated subunits resulting from autoproteolysis at high forces is thought to protect the integrity of the cell membrane when the cell is exposed to high mechanical stresses (75). In other SEAL domains, the role of this proteolysis is not well understood. For example, Notch proteolysis is required for function in in vivo fly experiments; yet, it is not required in in vitro mammalian cell experiments (76, 77). In dystroglycan, mutation of the autoproteolysis site S654A, which eliminates the catalytic hydroxyl group, leads to muscular dystrophy in mice and minor increase in cleavage by MMPs. Interestingly, many other disease-associated mutations, including C669F and I593D, cause deficient autoproteolysis while increasing MMP cleavage. The effects that modulating proteolysis has on cellular dynamics suggests it may play an important role in cells, though this needs to be further studied.

Even less understood is the role of MMP cleavage in dystroglycan function. Similar to Notch, the cleavage may be important to release the intracellular domain to affect transcription, as suggested by Azuara-Medina et al. (32). They and others suggest that dystroglycan plays an important role in cellular stress response though it is uncertain what exact role the intracellular domain of dystroglycan plays in this (33). Cleavage of dystroglycan is known to be important for brain plasticity where neuronal signals affect dystroglycan cleavage (31, 78). However, the physiological role MMP cleavage plays in muscle cells is unclear. As shown here, dystroglycan cleavage allows cells to migrate more freely, so it may be important for satellite cell response to muscle injury. It is also possible that dystroglycan cleavage acts a signaling pathway for cellular stress if there is too much strain on the DGC. Nevertheless, the pathogenic role of MMP cleavage in a variety of diseases is well known and here we show how it may play a role in dystroglycanopathies by altering cell mechanics.

An emerging theme is that SEAL domains play a role in communicating mechanical stimuli.. Indeed many of these proteins are involved in mechanotransduction events including ECM binding and cell-cell adhesion. Here we demonstrate that dystroglycan mutations decrease forces that cells exert on their surrounding tissues and affect cellular migration, most likely through crosstalk with other mechanosensitive complexes like integrin based focal adhesions which has been suggested in other studies (2, 79, 80). In this way, in addition to the possible mechanosignaling role discussed above, dystroglycan plays an important role in maintaining the integrity of the cell membranes in not only muscle cells but also likely in cancer where dystroglycan mutations and the increase in cellular migration is associated with metastasis (16, 81). Still, other SEALs require force application for less common functions. One of the more recent examples is ACE2 where force application causes ectodomain shedding which is thought to then bind to immune cells and cause a change in their response (82). It may even have an unknown role in blood pressure regulation through the renin-angiotensin-aldosterone system. Yet ACE2 shedding is most well known because of its role of viral entry, including SARS-CoV-2 virus which binds to ACE2 in order to enter the cell via regulated intramembrane proteolysis (RIP) (83). Furthermore, the recently discovered bacterial SEAL in *Bacillus subtilis* RsgI responds to membrane damage via RIP (44). Here the RsgI ectodomain binds to the damaged site, pulling on the SEAL domain which activates a damage response pathway. Protocadherin-15 is involved in mechanosensing during hearing and Notch detects forces associated with intercellular contact. Therefore, SEALs are involved in many cellular processes – both traditional mechanical pathways and not – and it would be interesting to compare the propensity of these domains to withstand forces using force spectroscopy..

Finally, this study shows that one possible therapeutic avenue to treat dystroglycanopathies would be inhibition of MMP cleavage. While broad spectrum MMP inhibitors had promising preclinical testing, all failed during Phase III study due to low efficacy, possibly because of their broad action (84–87). Therefore, a more specific inhibitor, such as a small molecule or nanobody, would need to be developed. The high-throughput techniques shown here, including flow cytometry readout of MMP cleavage and cellular forces could act as good screening tools for these inhibitors.

## SUPPLEMENTAL INFORMATION

**Figure S1.**
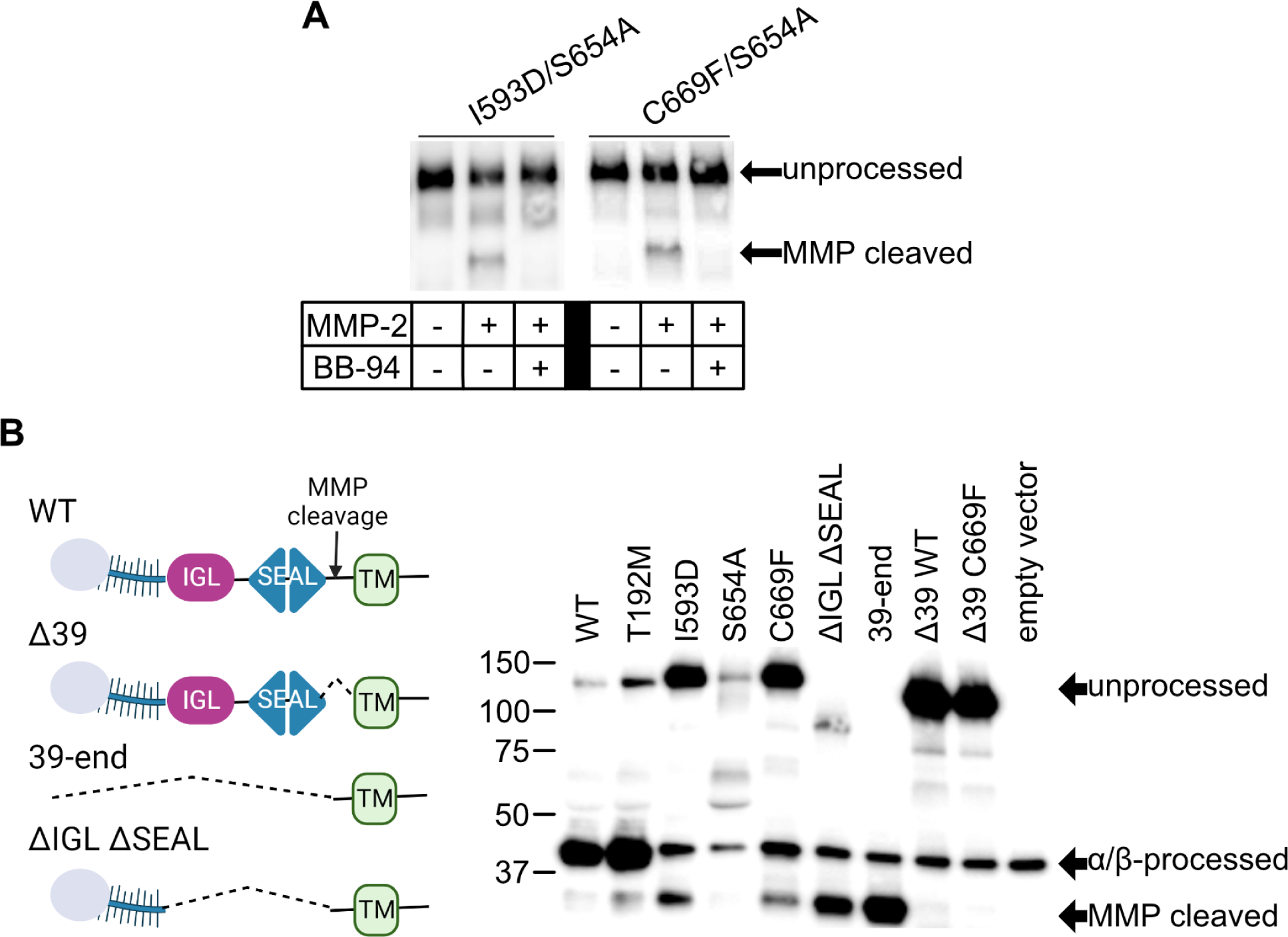
Dystroglycan cleavage products match expected size. **A)** Anti-HA Western blot of dystroglycan with double mutations exposed to 0.73 ug/mL +/− 40 uM BB-94. **B)** (left) Schematics of constructs used for Western blotting where dotted carat represents deleted sections. (right) Anti-β-dystroglycan Western blot of full-length dystroglycan overexpressed in Cos7 cells.

**Figure S2.**
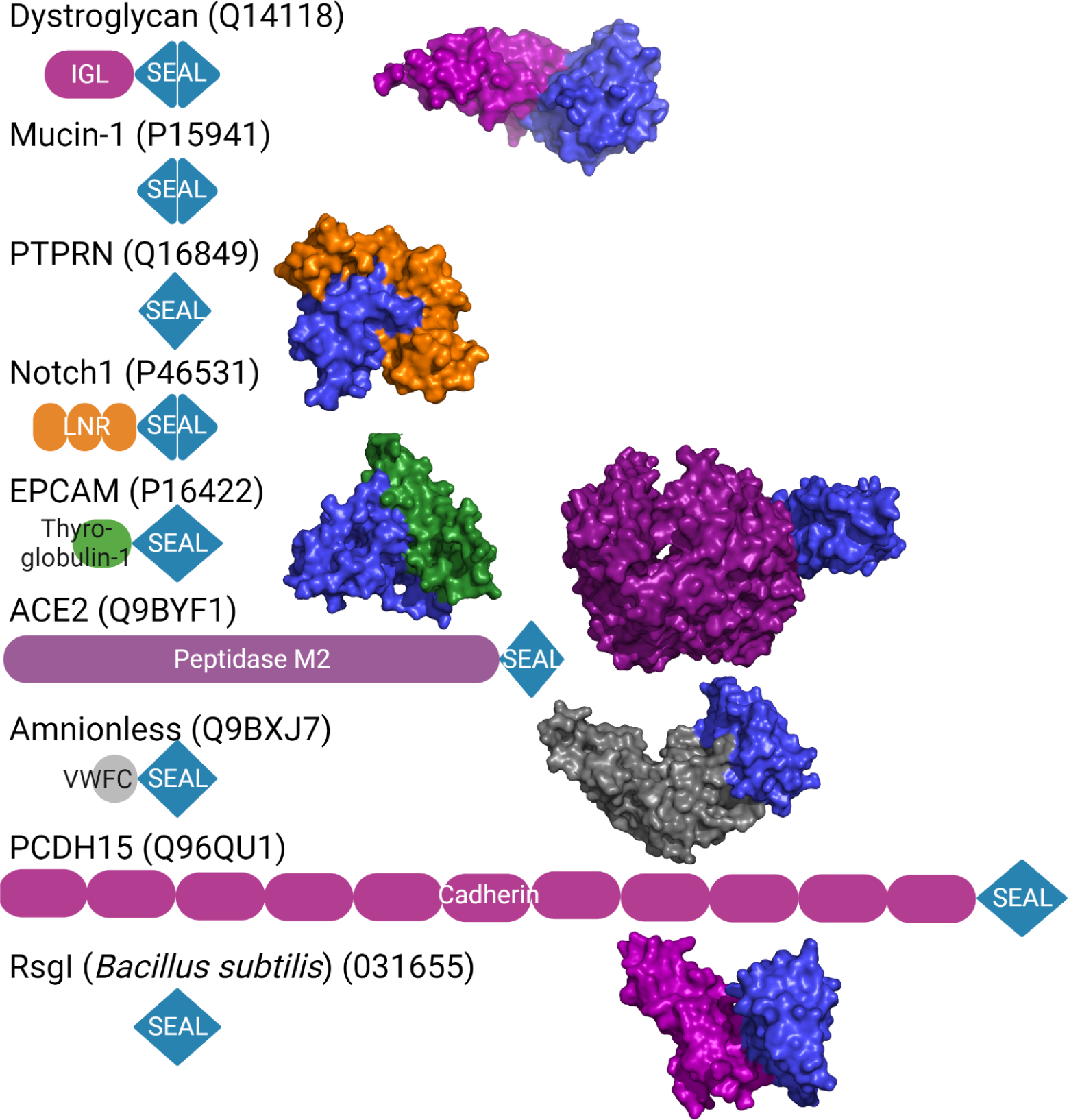
Comparison of SEAL neighboring domains. Schematics of SEAL domain-containing proteins with published crystal structure. Proteins with neighboring domains are shown in surface plot with SEAL domain in blue. Uniprot codes for proteins are shown in the parentheses.

**Figure S3.**
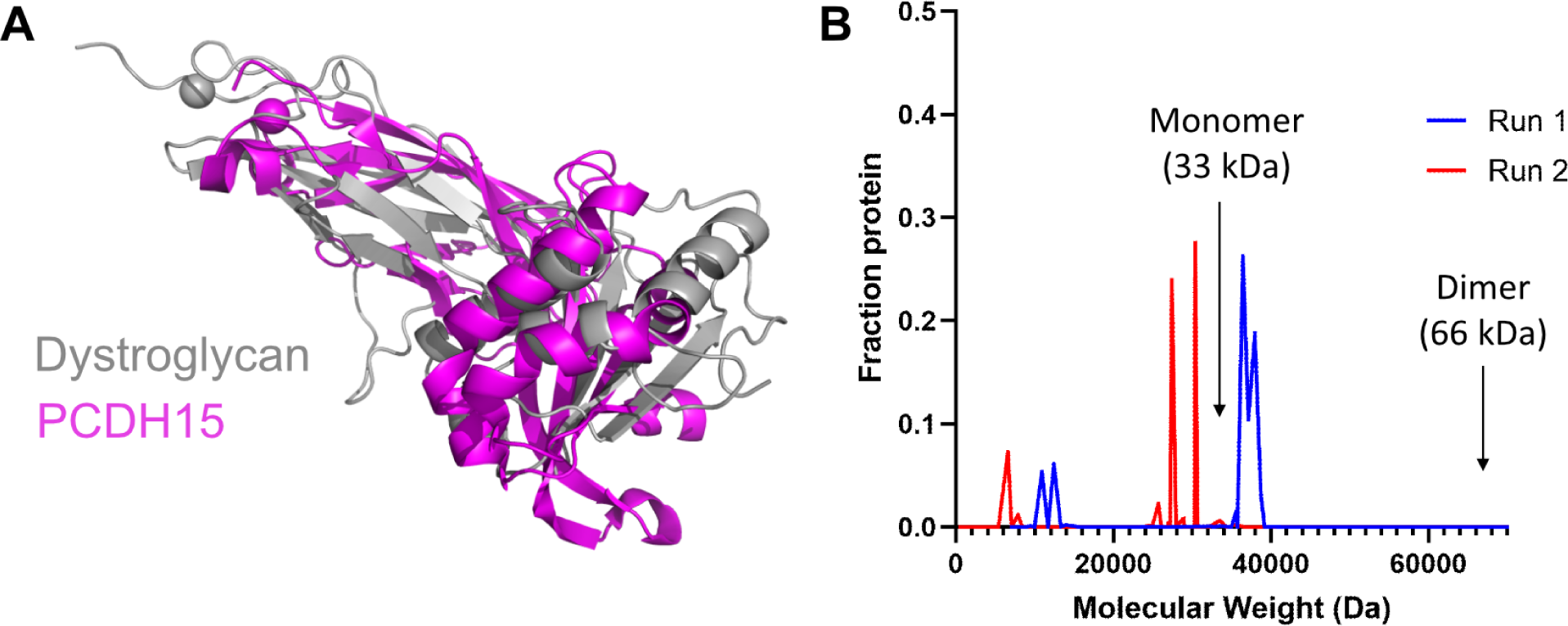
Dystroglycan is a monomer in solution. **A)** Dystroglycan (gray) and PCDH15 (magenta; PDB: 6BXZ) alignment. **B)** Calculated molecular weight by AUC for two biological replicates.

**Figure S4.**
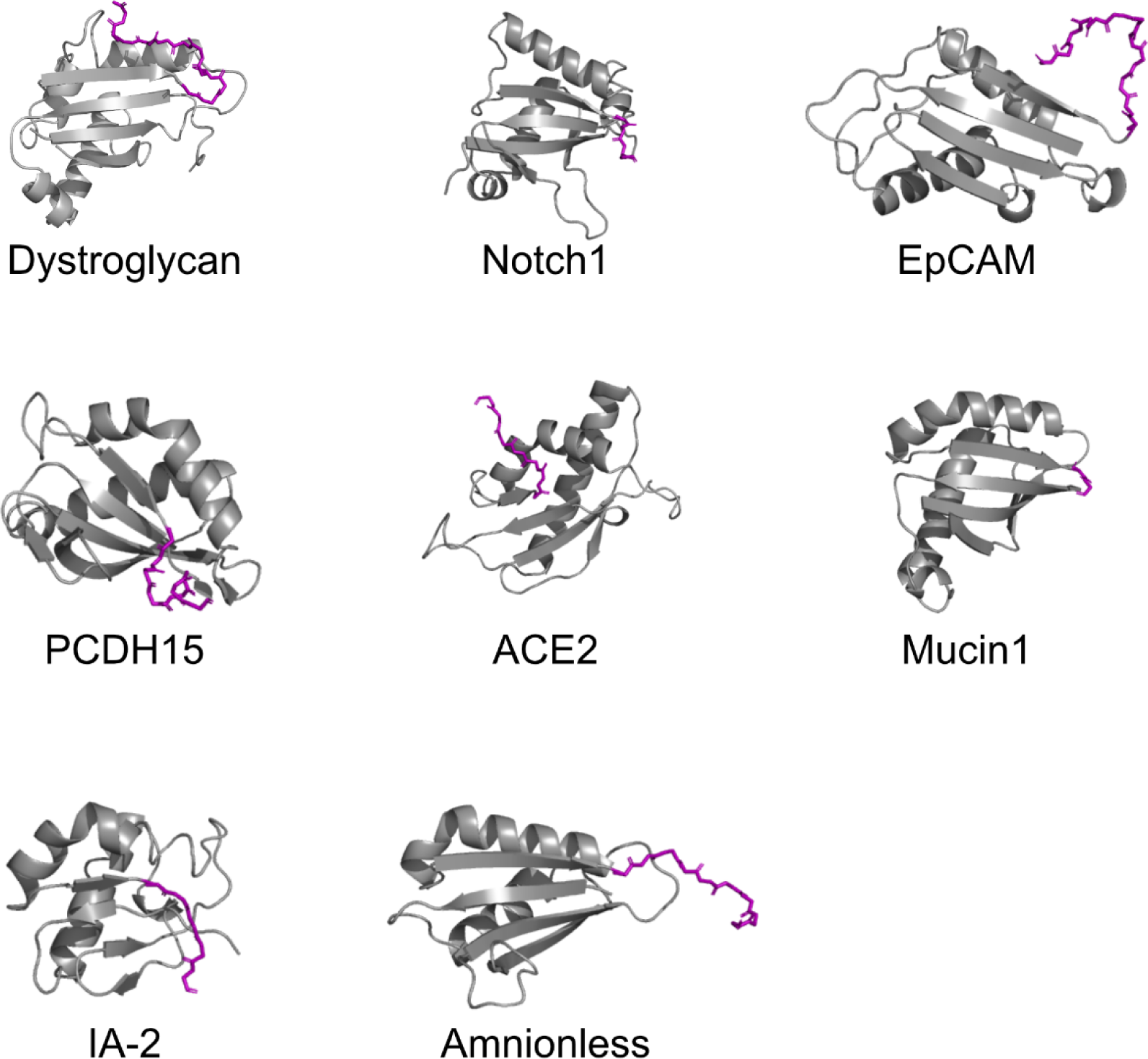
Comparison of SEAL C-termini. SEAL domains with C-terminus shown in purple. Only the C-terminus of dystroglycan interacts so closely with the SEAL domain.

**Figure S5.**
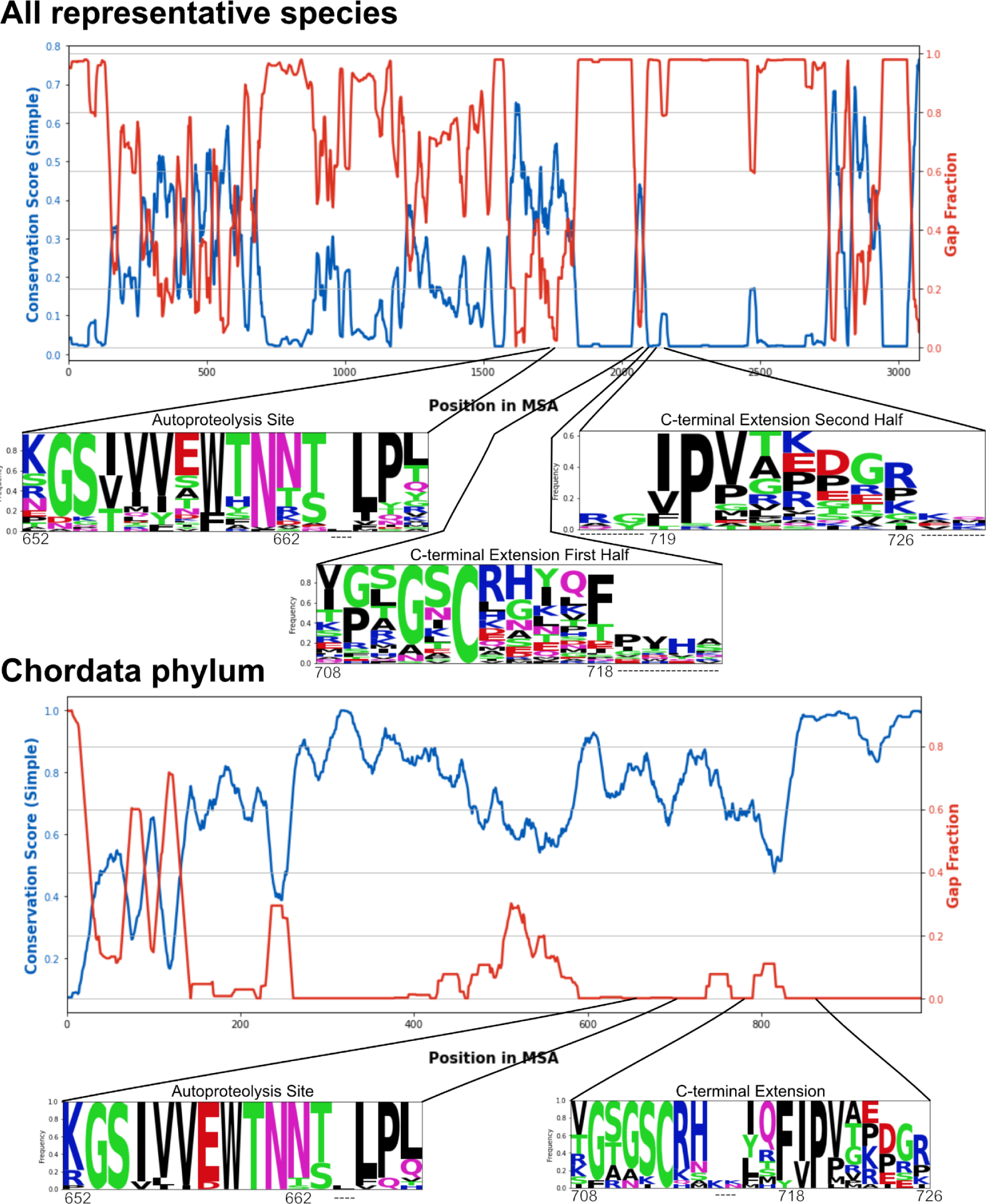
Dystroglycan autoproteolysis site and SEAL C-terminal extension are highly conserved. Conservation score and gap function for alignment of representative species from all phylums with dystroglycan (top) or just the Chordata phylum (bottom). Logo plots show the conservation for the autoproteolysis site and SEAL C-terminal extension with human dystroglycan amino acid numbers shown below. —-denotes that the amino acid is not found in human dystroglycan.

**Figure S6.**
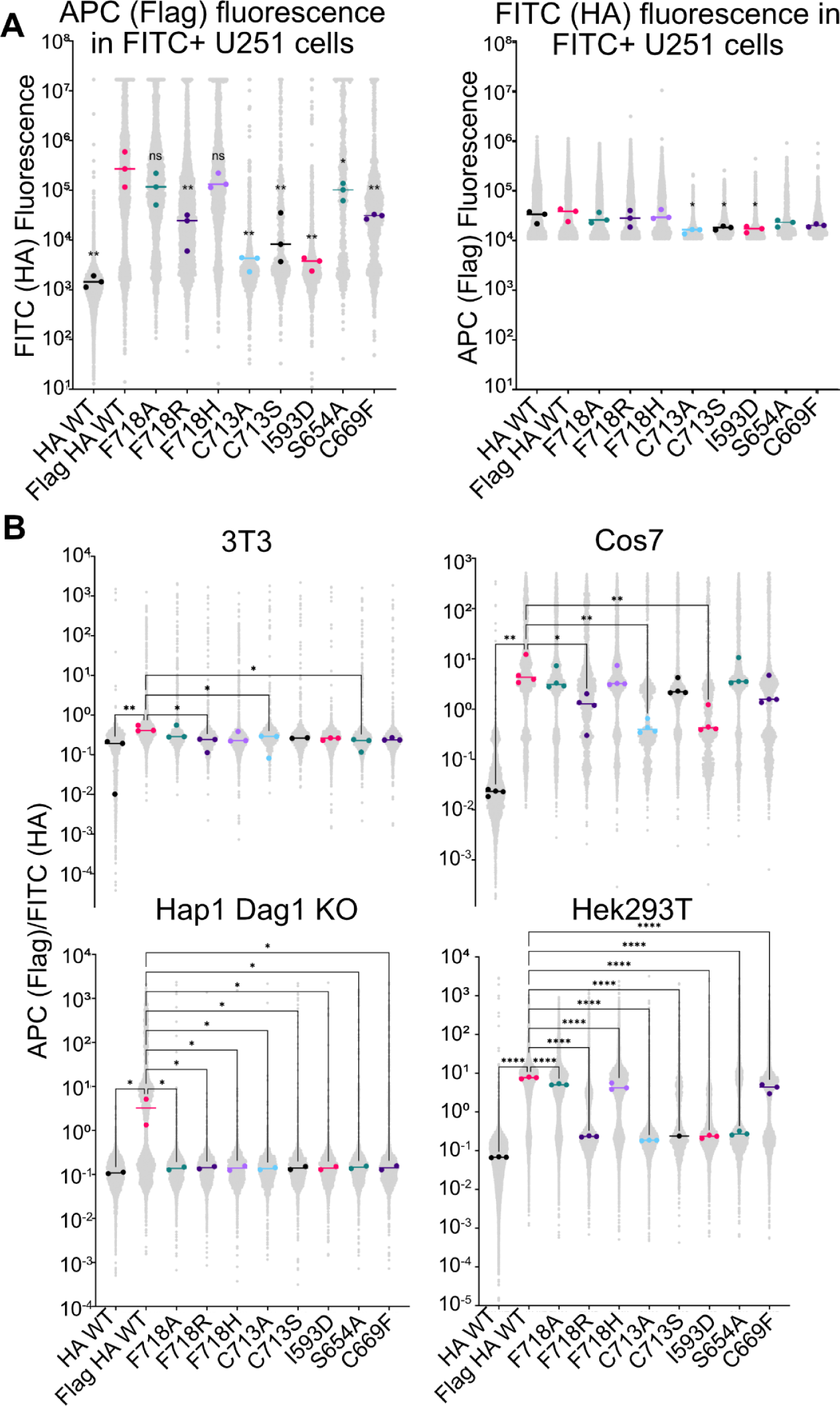
Dystroglycan is increasingly proteolyzed by structure-guided and muscular dystrophy-associated mutations. **A)** APC (left) and HA (right) individual values in FITC+ U251 cells. **B)**APC/FITC ratio for structure-guided and muscular dystrophy-associated mutations in FITC+ cells. Large dots represent median for individual days and line represents overall median. Significance was calculated using a one-way ANOVA with follow-up Dunnett’s multiple comparison test to Flag HA WT. * = p<0.05, ** = p<0.01, *** = p<0.001, **** = p<0.0001.

**Figure S7.**
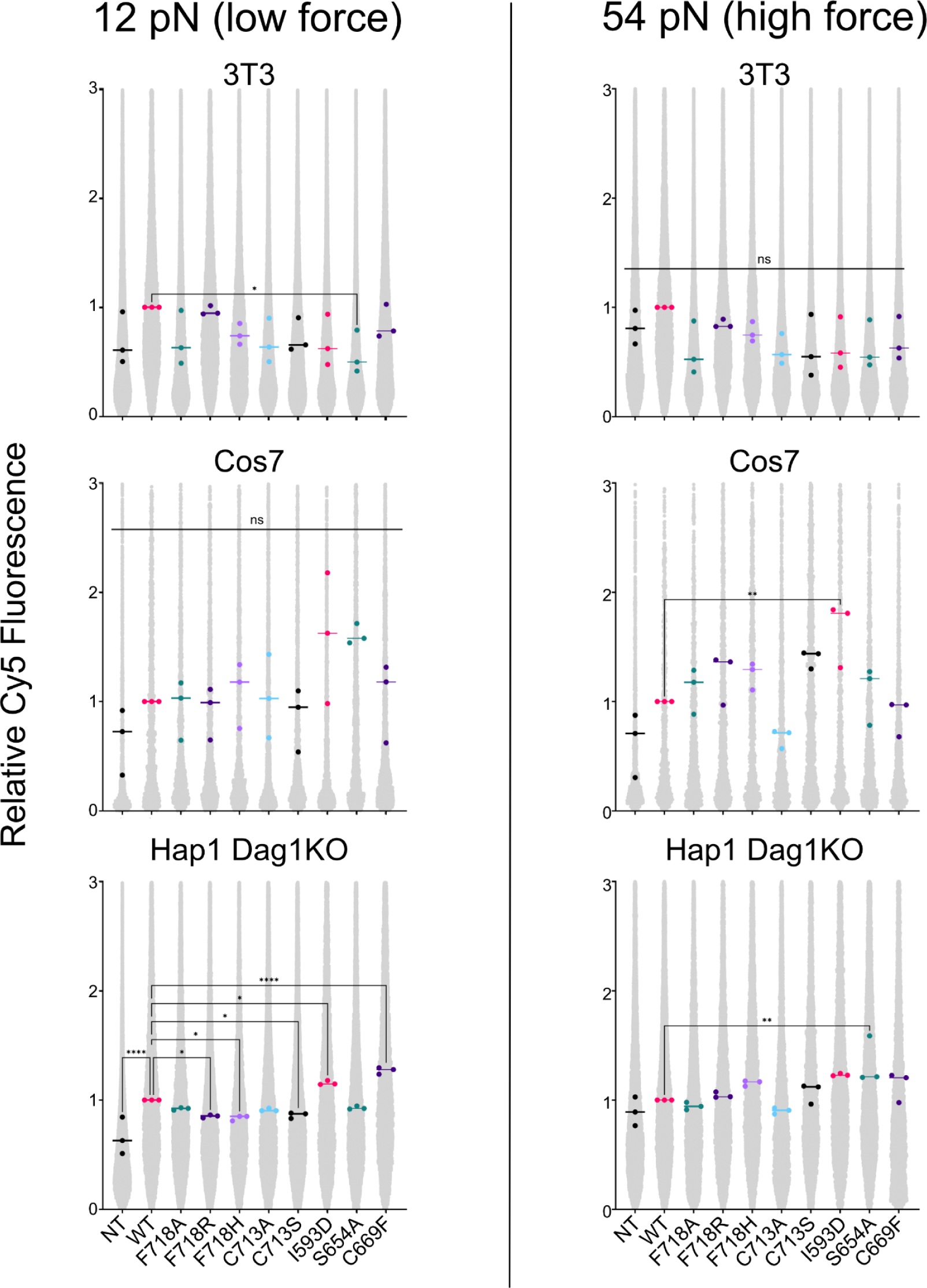
Dystroglycan cellular mechanics are altered by mutation. Relative Cy5 fluorescence compared to the median of WT for that day. Large dots represent median for individual day and line represents overall median. Significance was calculated via one way ANOVA with follow-up Dunnett’s multiple comparisons test to WT. NT = nontransfected, * = p<0.05, ** = p<0.01, *** = p<0.001, **** = p<0.0001.

**Figure S7A.**
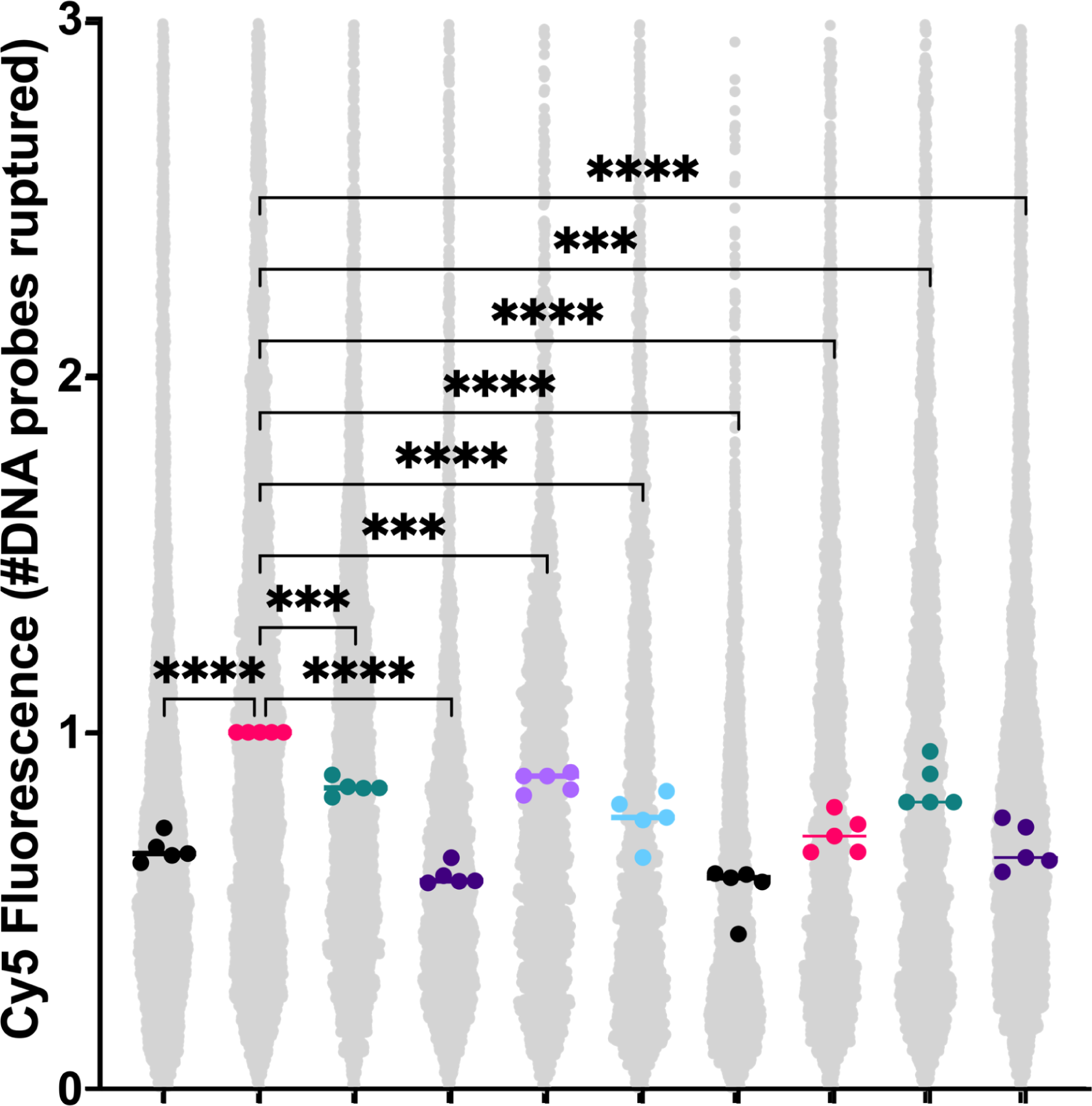
Dystroglycan cellular mechanics are altered by mutation. 12pN U251 cells from Figure 5B. Relative Cy5 fluorescence compared to the median of WT for that day. Large dots represent median for individual day and line represents overall median. Significance was calculated via one way ANOVA with follow-up Dunnett’s multiple comparisons test to WT. NT = nontransfected, * = p<0.05, ** = p<0.01, *** = p<0.001, **** = p<0.0001.

**Figure S8.**
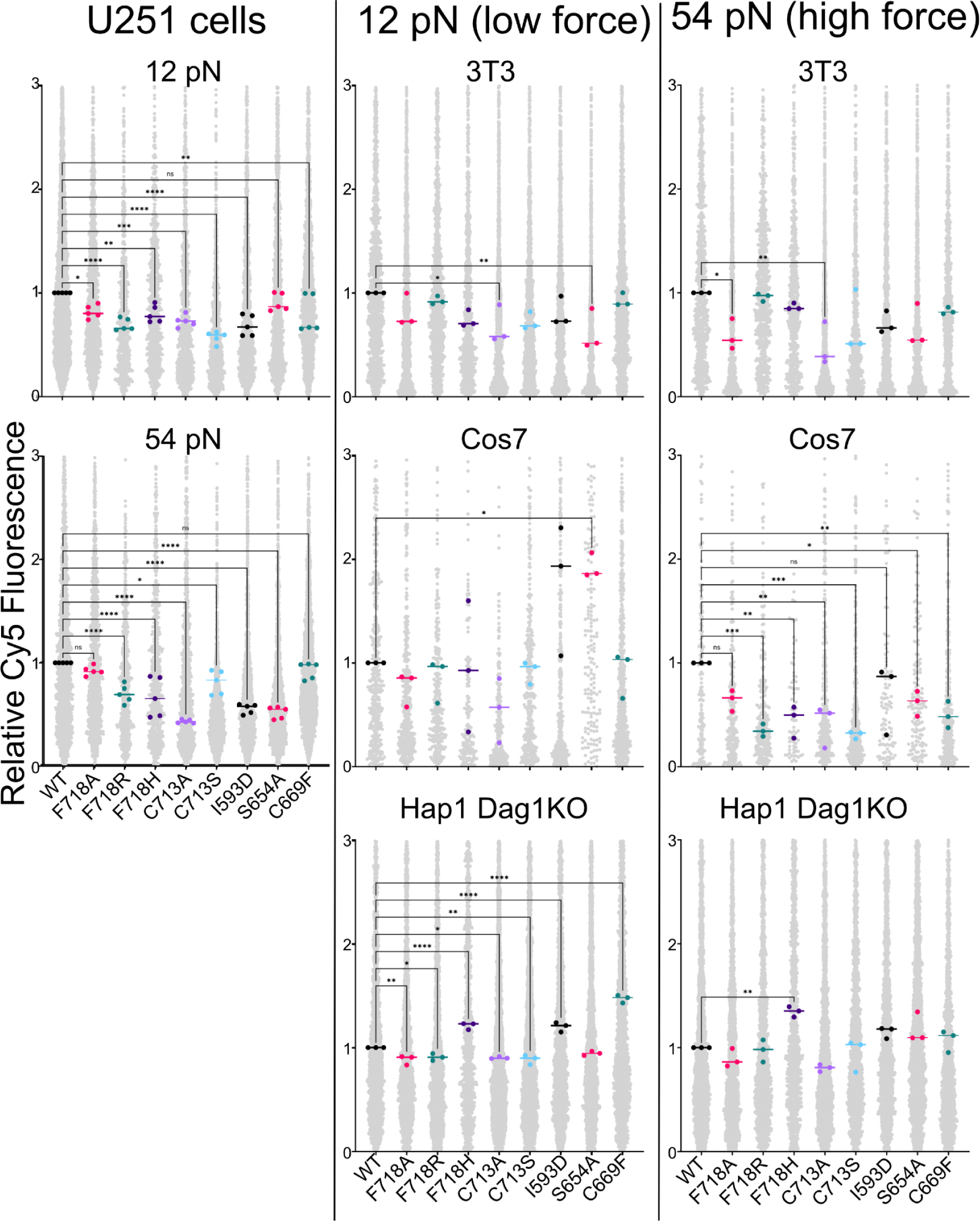
Cellular mechanics are similarly altered in GFP positive cells. Relative Cy5 fluorescence of GFP positive cells only compared to the median of WT for that day. Large dots represent median for individual day and line represents overall median. Significance was calculated via one way ANOVA with follow-up Dunnett’s multiple comparisons test to WT. NT = nontransfected, * = p<0.05, ** = p<0.01, *** = p<0.001, **** = p<0.0001.

**Figure S9.**
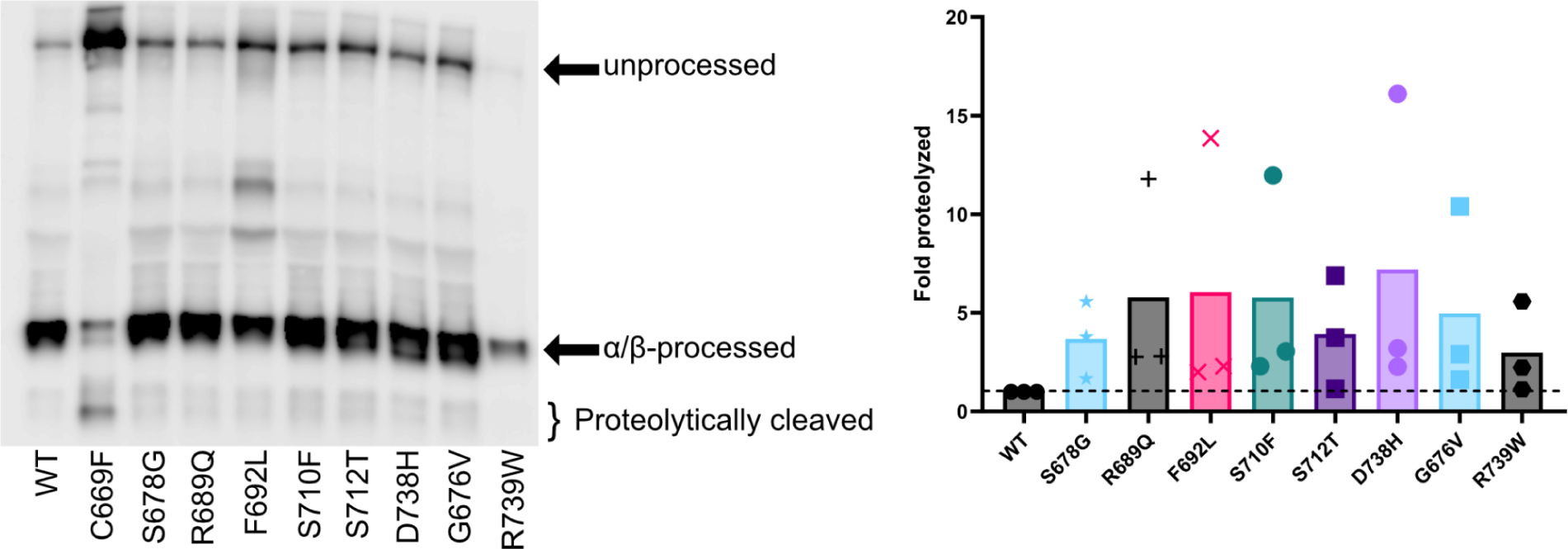
Cancer-associated mutations of dystroglycan increase MMP cleavage. Representative blot (left) and quantification (right) of proteolysis of anti-β-dystroglycan Western blot of overexpressed full-length dystroglycan in Cos7 cells.

**Figure S10.**
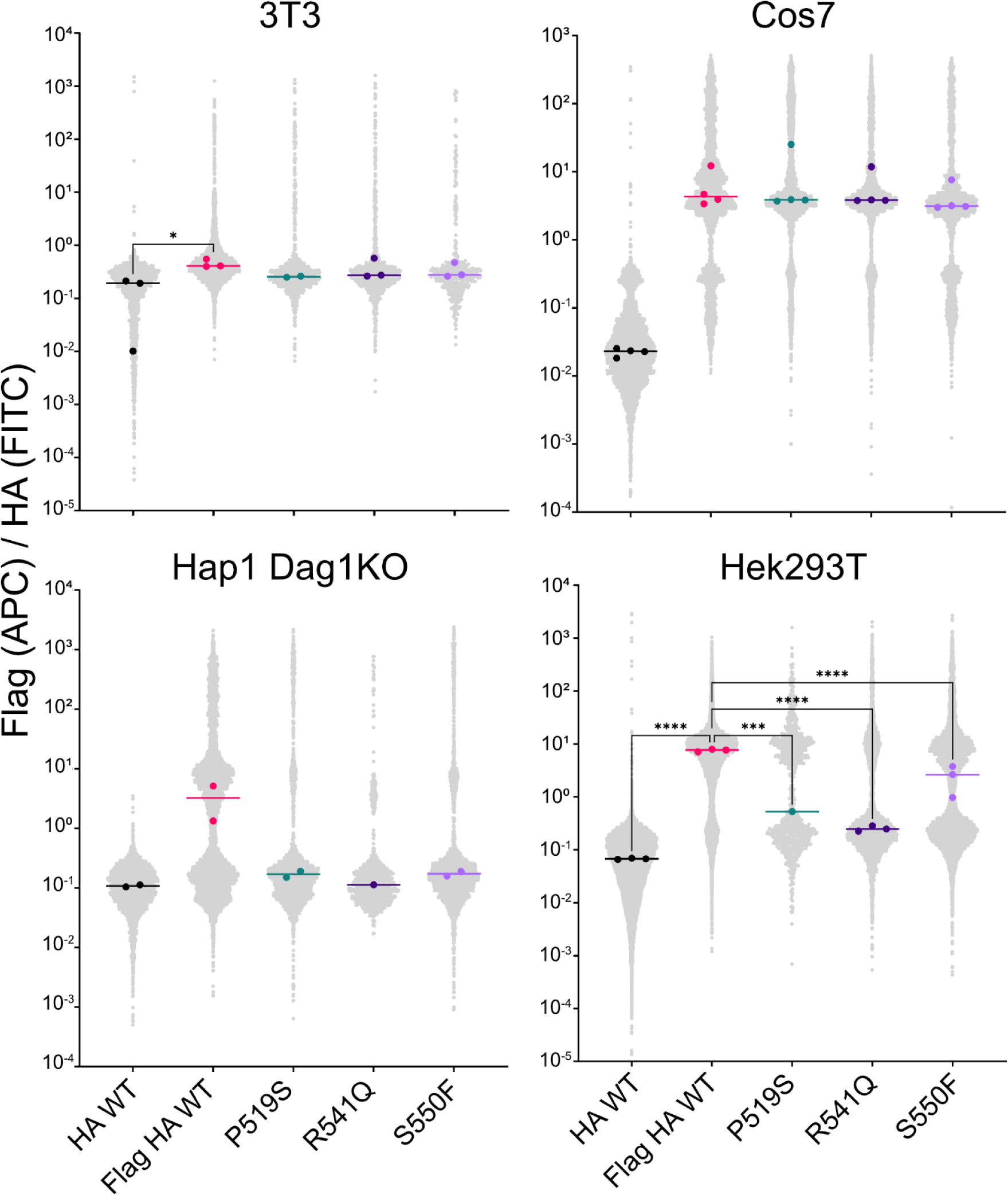
Dystroglycan proteolysis is increased in multiple cell lines as read out by flow cytometry. APC/FITC ratio for cancer associated mutations in FITC+ cells. Large dots represent median for individual days and line represents overall median. Significance was calculated using a one-way ANOVA with follow-up Dunnett’s multiple comparison test to Flag HA WT. * = p<0.05, ** = p<0.01, *** = p<0.001, **** = p<0.0001.

**Figure S11.**
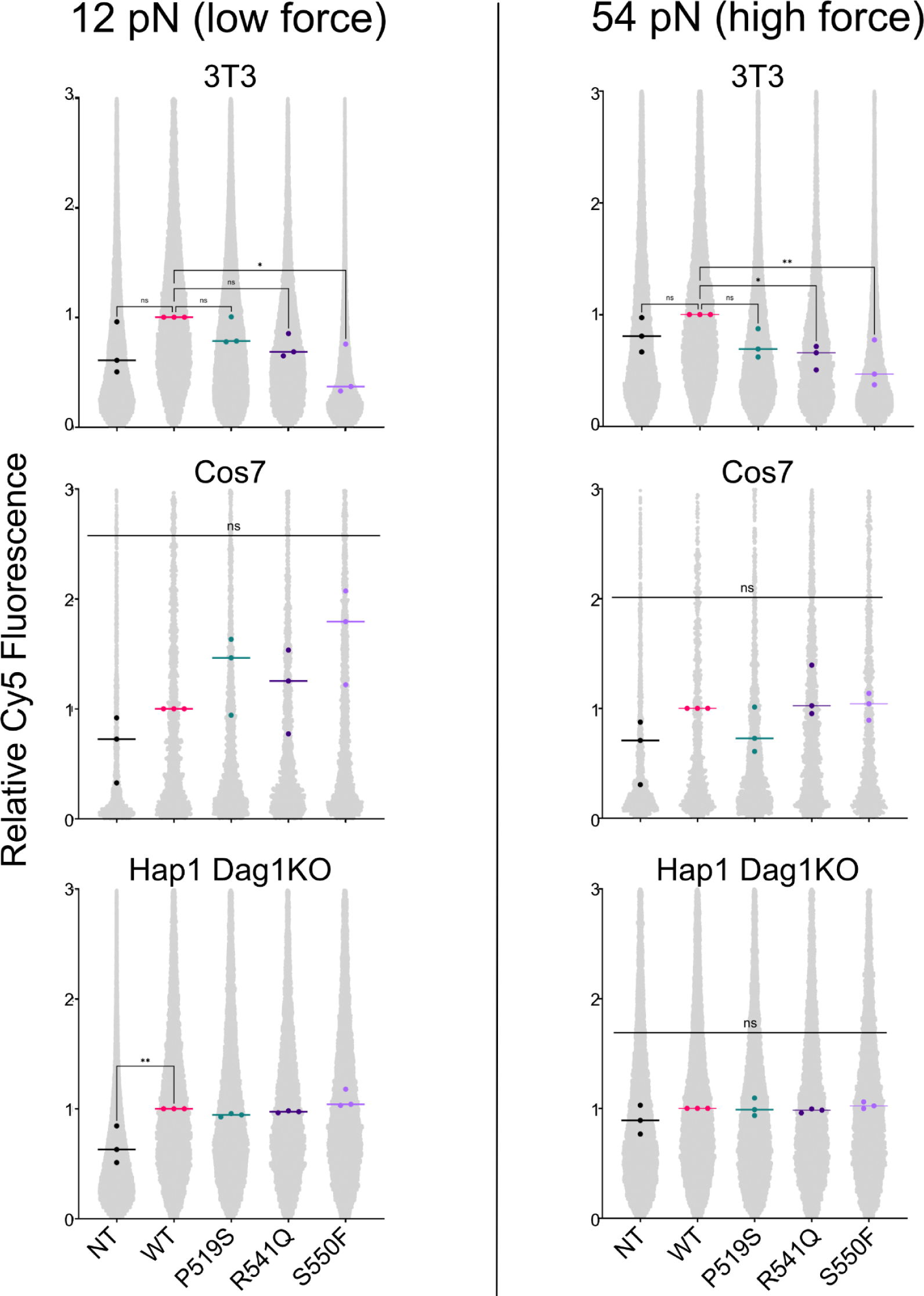
Dystroglycan cellular mechanics are altered by mutation. Relative Cy5 fluorescence compared to the median of WT for that day. Large dots represent median for individual day and line represents overall median. Significance was calculated via one way ANOVA with follow-up Dunnett’s multiple comparisons test to WT. * = p<0.05, ** = p<0.01, *** = p<0.001, **** = p<0.0001.

**Figure S12.**
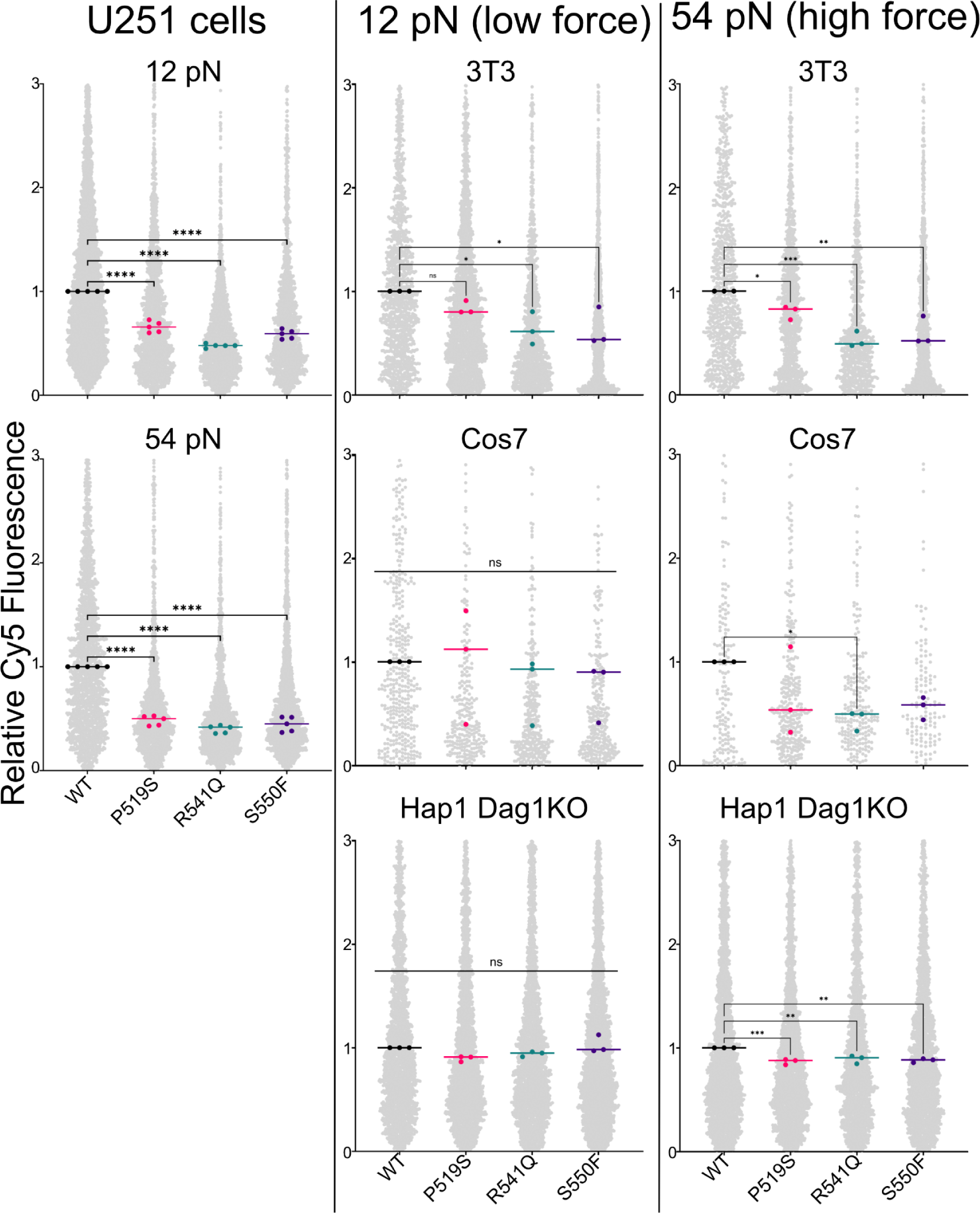
Cellular mechanics are similarly altered in GFP positive cells. Relative Cy5 fluorescence of GFP positive cells only compared to the median of WT for that day. Large dots represent median for individual day and line represents overall median. Significance was calculated via one way ANOVA with follow-up Dunnett’s multiple comparisons test to WT. NT = nontransfected, * = p<0.05, ** = p<0.01, *** = p<0.001, **** = p<0.0001.

**Figure S13.**
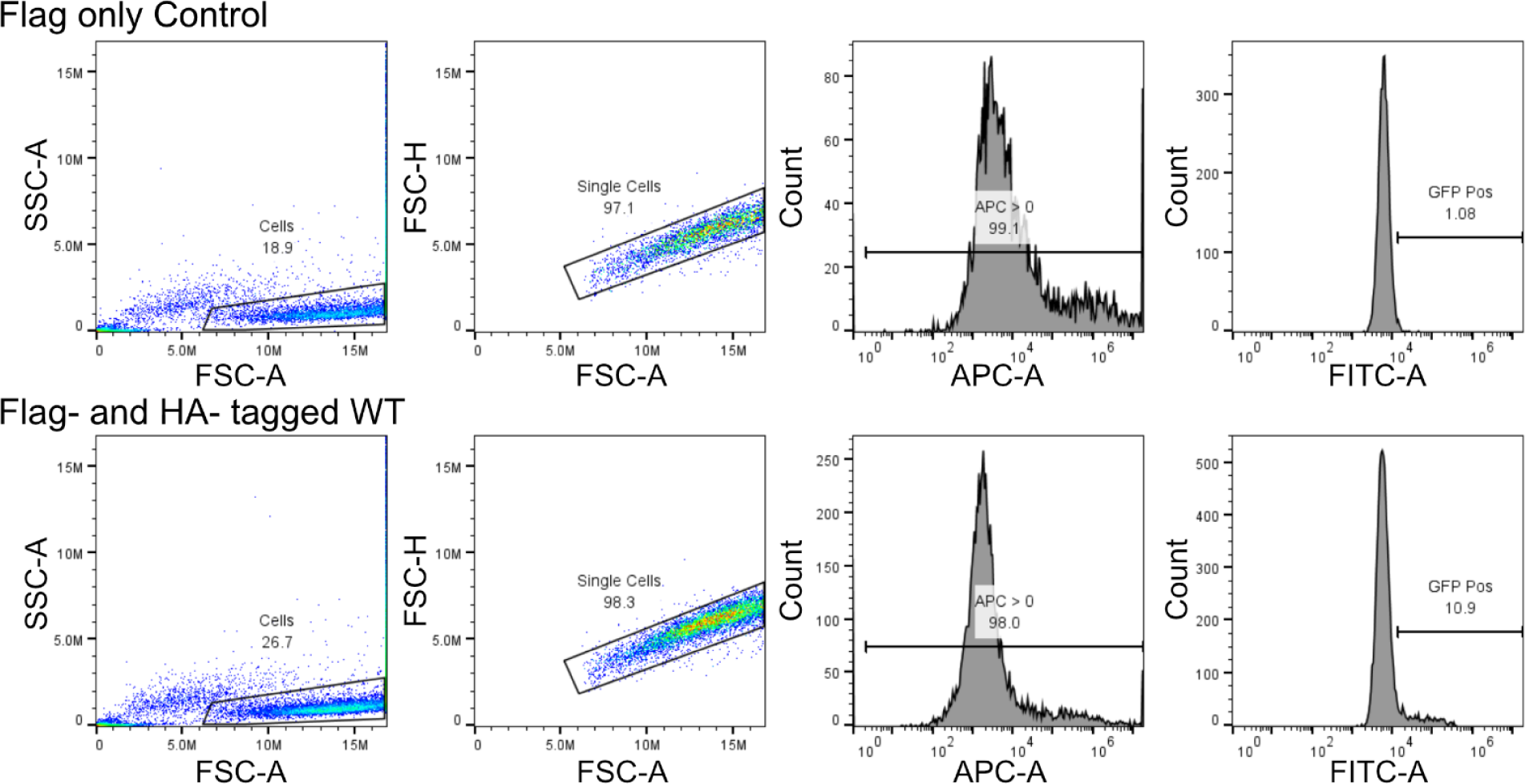
Gating strategy for flow proteolysis experiments.

**Figure S14.**
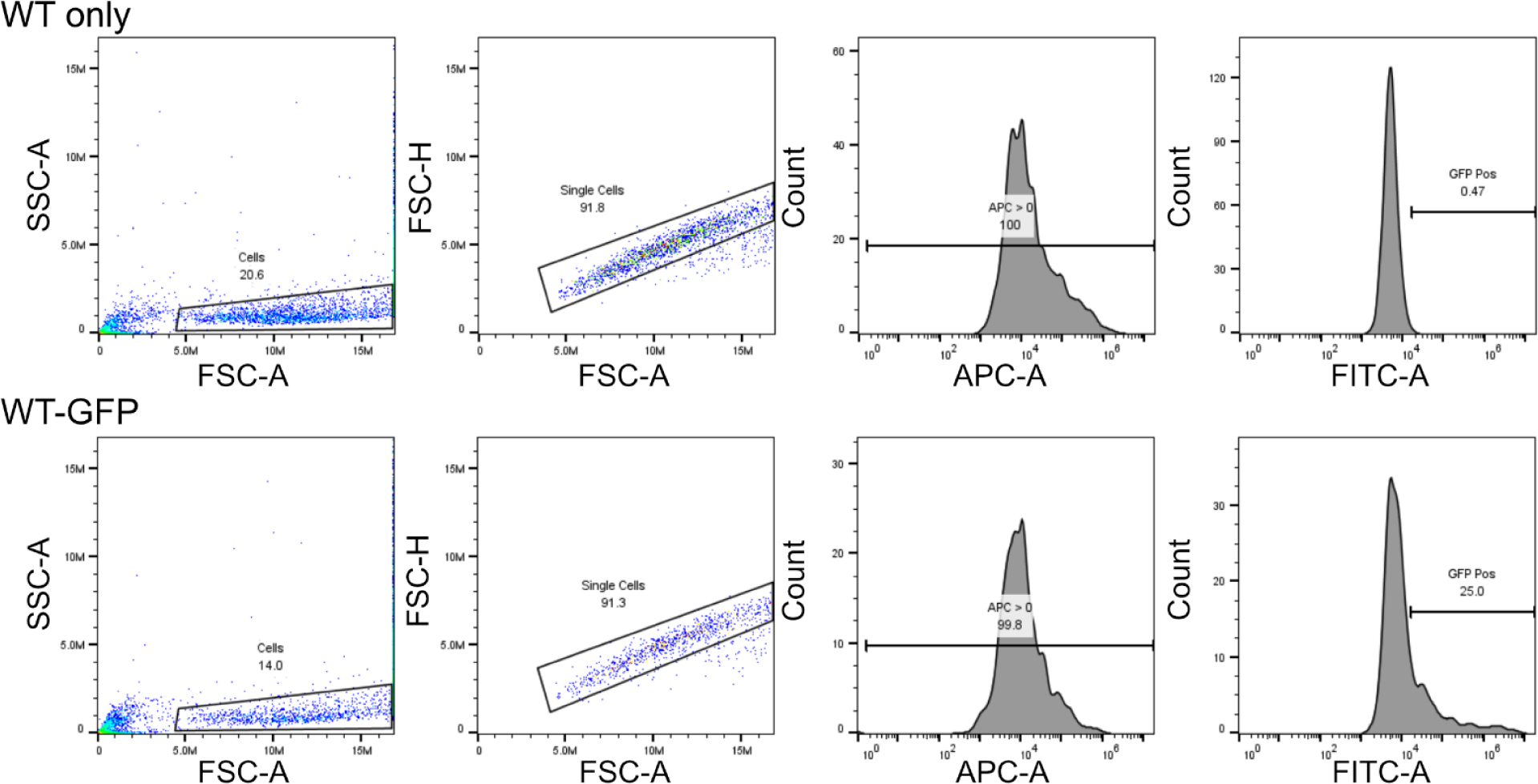
Gating strategy for RAD TGT experiments.

**Table S1.** Membrane-adjacent SEAL doamin comparison to dystroglycan using predicted structures from the AlphaFold Protein Structure Database (88).

| Protein | Uniprot Accession Number | SEAL RMSD with dystroglycan proteolytic domain (Å) | Neighboring domain | Buried surface area between neighboring domain and SEAL (Å<sup>2</sup>) |
| --- | --- | --- | --- | --- |
| Dag proteolytic domain | Q14118 |  | IGL | 568.6 |
| Sperm protein | Q07929 | 5.026 | EGF-like | 263.4 |
| Enterokinase | P98073 | 4.815 | None | 88.2 |
| Agrin | O00468 | 4.715 | None | 401.3 |
| Alpha-sarcoglycan | Q16586 | 3.134 | Cadherin-like | 615.8 |
| CDHR2 | Q9BYE9 | 5.416 | Cadherin | 510.3 |
| PTPRR | Q15256 | 1.393 | Alpha helix | 647.2 |
| Collectrin | Q9HBJ8 | 3.236 | Alpha helix | 260 |
| CD34 | P28906 | 5.362 | None | No interface predicted |
| KIAA0391 | Q5VV43 | 1.181 | PKD | 310.5 |
| NUP54 | Q7Z3B4 | 1.521 | None | 100.1 |
| DE-cadherin | Q24298 | 3.952 | Cadherin | 432.8 |

**Table S2.**
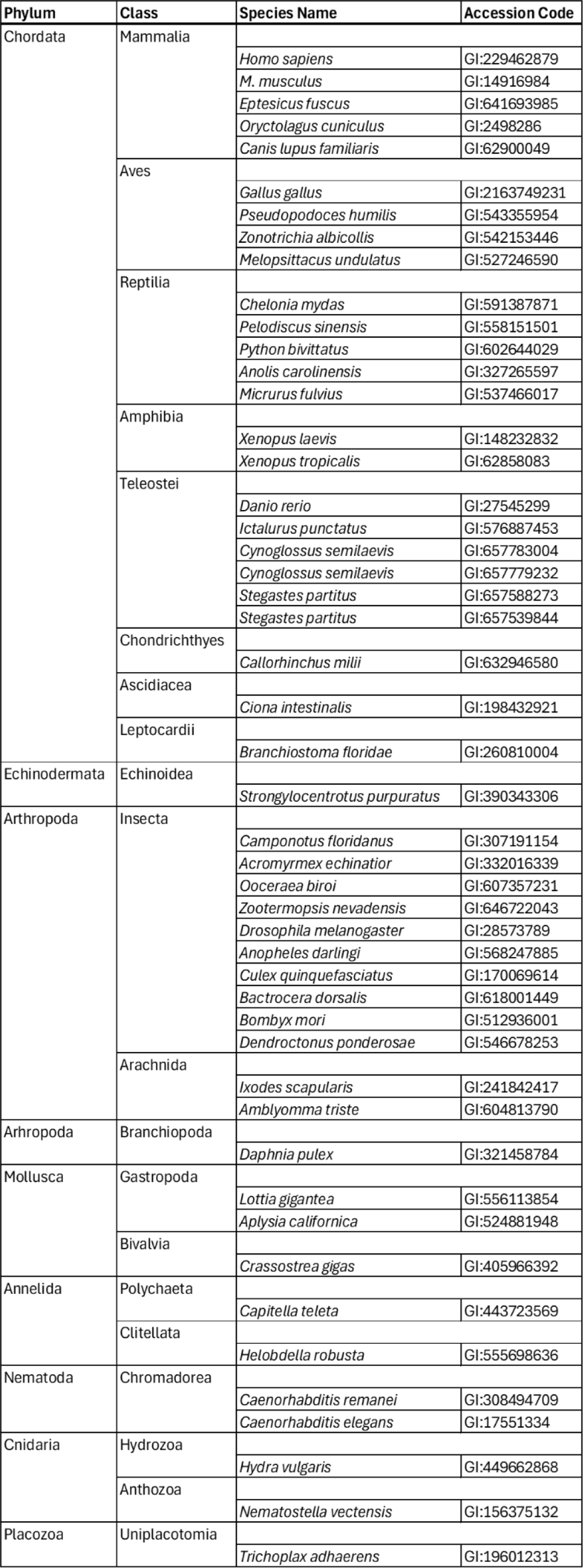
Species used for evolution analysis.

| Phylum | Class | Species Name | Accession Code |
| --- | --- | --- | --- |
| Chordata | Mammalia |  |  |
|  |  | *Homo sapiens* | GI:229462879 |
|  |  | *M. musculus* | GI:14916984 |
|  |  | *Eptesicus fuscus* | GI:641693985 |
|  |  | *Oryctolagus cuniculus* | GI:2498286 |
|  |  | *Canis lupus familiaris* | GI:62900049 |
|  | Aves |  |  |
|  |  | *Gallus gallus* | GI:2163749231 |
|  |  | *Pseudopodoces humilis* | GI:543355954 |
|  |  | *Zonotrichia albicollis* | GI:542153446 |
|  |  | *Melopsittacus undulatus* | GI:527246590 |
|  | Reptilia |  |  |
|  |  | *Chelonia mydas* | GI:591387871 |
|  |  | *Pelodiscus sinensis* | GI:558151501 |
|  |  | *Python bivittatus* | GI:602644029 |
|  |  | *Anolis carolinensis* | GI:327265597 |
|  |  | *Micrurus fulvius* | GI:537466017 |
|  | Amphibia |  |  |
|  |  | *Xenopus laevis* | GI:148232832 |
|  |  | *Xenopus tropicalis* | GI:62858083 |
|  | Teleostei |  |  |
|  |  | *Danio rerio* | GI:27545299 |
|  |  | *Ictalurus punctatus* | GI:576887453 |
|  |  | *Cynoglossus semilaevis* | GI:657783004 |
|  |  | *Cynoglossus semilaevis* | GI:657779232 |
|  |  | *Stegastes partitus* | GI:657588273 |
|  |  | *Stegastes partitus* | GI:657539844 |
|  | Chondrichthyes |  |  |
|  |  | *Callorhynchus milii* | GI:632946580 |
|  | Ascidiacea |  |  |
|  |  | *Ciona intestinalis* | GI:198432921 |
|  | Leptocardii |  |  |
|  |  | *Branchiostoma floridae* | GI:260810004 |
| Echinodermata | Echinoidea |  |  |
|  |  | *Strongylocentrotus purpuratus* | GI:390343306 |
| Arthropoda | Insecta |  |  |
|  |  | *Camponotus floridanus* | GI:307191154 |
|  |  | *Acromyrmex echinator* | GI:332016339 |
|  |  | *Ooceraea biroi* | GI:607357231 |
|  |  | *Zootermopsis nevadensis* | GI:646722043 |
|  |  | *Drosophila melanogaster* | GI:28573789 |
|  |  | *Anopheles darlingi* | GI:568247885 |
|  |  | *Culex quinquefasciatus* | GI:170069614 |
|  |  | *Bactrocera dorsalis* | GI:618001449 |
|  |  | *Bombyx mori* | GI:512936001 |
|  |  | *Dendroctonus ponderosae* | GI:546678253 |
|  | Arachnida |  |  |
|  |  | *Ixodes scapularis* | GI:241842417 |
|  |  | *Amblyomma triste* | GI:604813790 |
| Arthropoda | Branchiopoda |  |  |
|  |  | *Daphnia pulex* | GI:321458784 |
| Mollusca | Gastropoda |  |  |
|  |  | *Lottia gigantea* | GI:556113854 |
|  |  | *Aplysia californica* | GI:524881948 |
|  | Bivalvia |  |  |
|  |  | *Crassostrea gigas* | GI:405966392 |
| Annelida | Polychaeta |  |  |
|  |  | *Capitella teleta* | GI:443723569 |
|  | Clitellata |  |  |
|  |  | *Helobdella robusta* | GI:555698636 |
| Nematoda | Chromadorea |  |  |
|  |  | *Caenorhabditis remanei* | GI:308494709 |
|  |  | *Caenorhabditis elegans* | GI:17551334 |
| Cnidaria | Hydrozoa |  |  |
|  |  | *Hydra vulgaris* | GI:449662868 |
|  | Anthozoa |  |  |
|  |  | *Nematostella vectensis* | GI:156375132 |
| Placozoa | Uniplacotomia |  |  |
|  |  | *Trichoplax adhaerens* | GI:196012313 |

## MATERIALS AND METHODS

### Plasmid Creation

Full-length human dystroglycan cDNA in CMV-6 plasmid was obtained from Origene and used as a template for all of the cloning. Cloning was performed using In-Fusion HD Cloning Mix (Takara). Restriction enzymes were purchased from New England Biolabs (NEB). All electrophoresis supplies were purchased from Bio-Rad. All primers for cloning were purchased from Integrated DNA Technologies (IDT). All mutagenesis was done using instructions from In-Fusion HD Cloning Mix (Takara) unless noted below.

For secreted proteolysis domain: IGL + SEAL domain (amino acids 490-749) and SEAL domain (599–749) constructs were cloned into pcDNA3 vector containing an N-terminal Notch signal sequence and Flag tag and C-terminal Fc-fusion and HA tags. Site-directed mutagenesis was done using Pfu Turbo Polymerase (Agilent).

For Western blots: Full-length human dystroglycan was cloned into pcDNA5 vector.

For X-ray crystallography/AUC: IGL + SEAL domains (amino acids 491-748) construct was cloned into pTD68 vector containing N-terminal 6x His tag and Sumo tag.

For flow cytometry proteolysis assays: Full-length human dystroglycan was cloned into pcDNA5 vector. A Flag tag x3 was inserted between amino acids 489 and 490 and a HA tag was inserted between amino acids 724 and 725 using In-Fusion Cloning Mix (Takara).

For RAD TGT: Full-length human dystroglycan was cloned into pcDNA5 vector with C-terminal mNeonGreen.

### Mammalian Cell Culture

All media was supplemented with 10% fetal bovine serum (R&D), 100 units/mL penicillin (Sigma), and 100 units/mL streptomycin (Sigma). Cells were grown in Dulbecco’s Modified Eagle Medium (DMEM, Corning; U251, NIH-3T3, Hek293T, Cos7) or Iscove Modified Dulbecco Medium (IMDM, Gibco; Hap1 Dag1 KO). Cells were kept only up to 20 passages and were passaged prior to reaching 90% confluency.

Transfections were done in Opti-MEM (Gibco) with Lipofectamine 3000 (Invitrogen) according to manufacturer instructions.

COS-7 cells were a gift from Dr. Margaret Titus. U251 cells were a gift from Dr. David Odde. NIH-3T3 cells were a gift from Dr. Steven Blacklow. HEK293T cells were purchased from ATCC. HAP-1 DAG1 KO cells were purchased from Horizon.

### Western Blots

Cell lysates from 24-well plates were collected 48 hours post-transfection using RIPA lysis buffer (Boston Bio Products). Lysates were electrophoresed on a 4-20% SDS-PAGE gel in Tris/Glycine/SDS running buffer (Bio-Rad) supplemented with 2 mM sodium thioglycolate (Sigma). The protein was then transferred to a nitrocellulose membrane (Invitrogen) using a Genie Blotter (Idea Scientific) and blocked with 5% milk. The primary antibodies (details below) were diluted 1:1000 in Tris buffered saline (TBS, Thermo) supplemented with 0.1% Tween-20 (Sigma), 5% bovine serum albumin (BSA, Fisher), and 0.2% sodium azide (Sigma) and incubated overnight at 4°C. A goat-anti mouse HRP conjugated antibody (Invitrogen) was diluted 1:1000 in TBS with Tween-20, BSA, and sodium azide and incubated at room temperature for 1 hour. The membrane was then imaged using chemiluminescent buffer (Perkin Elmer Western Lightning Plus ECL) and an Amersham 600UV (GE) with support from the University of Minnesota Imaging Center. Unprocessed, processed, and cleaved bands were quantified using ImageJ.

Percent cleaved was calculated by dividing the sum of the cleaved bands by the sum of the unprocessed and processed bands. Fold change was calculated by dividing the percent cleaved for that sample by the percent cleaved of the WT for that blot. Fold change values were plotted using GraphPad Prism Software Version 9.4.1 where each dot represents a biological replicate and the bar represents the median.

Primary antibody for the secreted MMP cleavage experiments was anti-HA (Covance, MMS-101P). For the full-length dystroglycan blots, anti-β-dystroglycan (Leica, 43Dag1/8D5) was used.

### MMP Activation

Recombinant human MMP-2 (R&D) and MMP-9 (R&D) were activated using p-aminophenylmercuric acetate (APMA, Sigma) in MMP assay buffer (50 mM Tris (Fisher), 10 mM CaCl2 (Millipore), 150 mM NaCl (Fisher), 0.05% Brij-35 (w/v) (Millipore), pH 7.5) according to manufacturer instructions. MMP-2 was activated for 1 hour at 37°C, and MMP-9 was activated for 24 hours at 37 C.

### MMP Cleavage Assay

10 cm plates of Hek-293T cells were transiently transfected with dystroglycan IGL and SEAL domains. The conditioned media was collected 48 hours post-transfection. Conditioned media was bound to magnetic protein A beads (Invitrogen) and rotated for 1 hour at 4°C. The tube was then placed in a magnetic tube holder and supernatant was aspirated off. The beads were washed thoroughly with wash buffer (50 mM Tris, 150 mM NaCl, 0.1% NP-40 (VWR), pH 7.5). Beads were then split into tubes and MMP assay +/− activated MMP (final concentration 0.73 ug/mL unless noted otherwise) and +/− Batimastat (final concentration 40 uM) (BB94, VWR) as needed for that condition. Tubes were rotated for 1 hour for 37°C. Beads were then added to the magnetic tube holder and supernatant was aspirated off. Remaining protein was eluted using 2X SDS (Biorad) sample buffer with 100 mM DTT (Sigma), which was incubated for at least 5 minutes at room temperature. Samples were then run on a Western blot using anti-HA antibody as detailed above.

### Protein Expression and Purification

Sequence confirmed pTD68 vector with dystroglycan (amino acids 491-748) was transformed into BL21 (DE3) *E. coli* competent cells (Agilent). Cells were cultured in 1L LB broth at 37°C and induced at OD600 of 0.6 with 0.5 mM IPTG (isopropyl-d-1-thio-galactopyranoside) (Fisher) and grown overnight at 18°C. Cell were then pelleted by centrifuging at 4000 x g for 30 minutes at 4°C. Pellets were resuspended in 10 mL of lysis buffer (50 mM Tris, 250 mM NaCl, 1 mM EDTA (Fisher), pH 7.5) and pulse sonicated for 30 seconds 3 times with 2 minute rest on ice in between sonication rounds. Lysate was centrifuged at 24k x g for 30 minutes and supernatant discarded. To solubilized inclusion bodies, the pellet was resuspended in 10 mL solubilization buffer (50 mM Tris, 250 mM NaCl, 5 M Urea (Fisher), 1 mM EDTA, pH 7.5) and sonicated as before. This was again centrifuged at 24k x g for 30 minutes. The supernatant was batch bound to 2 mL HisPure Ni-NTA agarose beads (Thermo) for 1 hour at 4°C. After incubation, the mixture was added to a column and allowed to flow by gravity. Beads were washed with 20 mL each of wash buffer 1 (50 mM Tris, 500 mM NaCl, 3 M urea, pH 7.5) and wash buffer 2 (50 mM Tris, 200 mM NaCl, 100 mM imidazole (Fisher), 3 M urea, pH 7.5). Protein was eluted using 5 mL elution buffer (50 mM Tris, 200 mM NaCl, 250 mM imidazole, 1 M urea, pH 7.5).

To refold the protein, the Ni column eluant was mixed with an equal volume of refold buffer (50 mM Tris, 200 mM NaCl, 50 mM CaCl_2_ (Sigma), 2 mM cysteine (Fisher), 0.5 mM cystine (Fisher), pH 8.5) and placed in a 10 kDa snakeskin dialysis bag (Thermo). This was placed into 2L refold buffer and left at 4°C overnight. The following morning, the snakeskin was moved into 2L of fresh refold buffer and incubated at 4°C for at least 4 hours when it was once again placed into 2L of fresh refold buffer and incubated at 4°C overnight. For dialysis, the dialysis bag was placed into 2L dialysis buffer (50 mM Tris, 200 mM NaCl, 10 mM CaCl2, pH 7.5) and incubated at 4°C for at least 4 hours. 5 Units of ULP-1 per 1 L of *E. coli* expression was then added into the dialysis bag which was placed into 2L of fresh dialysis buffer and placed at 4°C overnight.

The contents of the dialysis bag was batch bound for at least 1 hour at 4°C with 2 mL HisPur Cobalt Resin (Thermo) to remove His-ULP-1 and His-SUMO that was cut from the dystroglycan. The mixture was added to a column and allowed to flow by gravity; the flow-through was collected as it contains the refolded dystroglycan. The column was washed with 10 mL dialysis buffer which was also collected and pooled with flow through. These were then concentrated using 3 kDa cut-off spin concentrator (Amicon Ultra-15 CentrifugalFilter Unit) to <2 mL in order to be purified via size exclusion column. Samples were purified using an ENrich SEC70 column (Bio-Rad) into filtered and degassed dialysis buffer. They were then concentrated to 7 mg/mL using spin concentrators.

### Crystallization, Data Collection, and Processing

Rigaku’s CrystalMation system was used for broad, sitting drop screening of dystroglycan crystallization. Initial crystals formed in 0.2 M lithium sulfate, 0.1 M Tris hydrochloride (pH 8.5), 30% (w/v) PEG 4000. Crystal refinement was done using 24-well hanging drop plates and siliconized thick glass cover slides (Hampton). Multiple conditions grew crystals with the best forming in a 1:1 ratio with the well condition of 0.1 M Tris (pH 8.3) (Hampton), 0.2 M lithium sulfate (Hampton), 20% PEG 4000 (Hampton). No cryoprotectant was used as this lowered the resolution during data collection. Crystals were looped, flash cooled in liquid nitrogen, and stored in liquid nitrogen until shipment to the Advanced Photon Source at the Argonne National Laboratory. Data was collected at APS Beamline 24 (NE-CAT) and processed using the HKL suite (89).

### Structural Prediction and Refinement

The dystroglycan structure could not be properly refined using homologous proteins including Notch1 NRR (PDB: 3ETO), PCDH15 (PDB: 6BXZ), and Mucin (PDB: 2ACM). We, therefore, used AlphaFold2^72^ to create a structural prediction of the same amino acids used for crystallization. Using the default settings on ColabFold^73^, we received 5 structural predictions which largely looked the same. The first prediction was used as a model for molecular replacement in PHENIX Software Version 1.19.2 (90). Remarkably, this worked to give an initial structure that could be used for further refinement. The electron density map was visualized using *Coot* Software Version 0.9.4.1 (91) which was also used for structure model correction. Refinement was done by alternating use of PHENIX auto.refine tool (with default settings) and visual correction in *Coot*. Structural validation was carried out in PDB-REDO (92) and MolProbity (93).

### Structural Analysis

Protein structures were visualized and images were created using PyMOL Software Version 2.5.7 (94). The superimposition function was used to align protein structures and calculate RMSD between two structures. Superimposition was always done using just the backbone structure to eliminate any bias from residue side chains.

### Flow-Based Dystroglycan Proteolysis

Cells were grown in a 48 well dish until approximately 90% confluent and transiently transfected as detailed above. 24 hours later, cells were tryspinized until dislodged from the plate when 250 uL of media supplemented with 10% FBS was added. Cells were transferred to a 1.5 mL microcentrifuge tube and centrifuged at 400 x g for 3 minutes. The supernatant was aspirated off and cells were washed with DPBS (Corning). Samples were resuspended in Optimem with a 1:1000 dilution of APC conjugated to anti-Flag (PerkinElmer) and 1:500 dilution Alexa Flour 488 conjugated to anti-HA (Invitrogen) and rotated at 4°C, covered from the light for 1 hour. Cells were spun down at 400 x g for 3 minutes and washed twice with DPBS. They were then resuspended in 250 uL of Optimem and placed on ice in the dark until run on the cytometer.

The cytometer used for all data collection was a BD Accuri C6 Plus Personal Flow Cytometer using BD Accuri C6 Plus (64-bit) Software Version 1.034.1. Prior to each experiment the cytometer was checked for proper function using BD CS&T RUO Beads (Fisher) and the quality control function on the software. Furthermore, to ensure specific binding of the fluorescent antibodies, each experiment included WT dystoglycan without any tags, WT dystroglycan with Flag tag only, and WT dystroglycan with HA tag only as controls. All samples were run until 200,000 events or 220 uL of volume was reached.

Samples were analyzed using FlowJo (64-bit) Software Version 10.8.1 (BD Life Sciences). Data was gated on single cells with fluorescent signal > 0 (Figure S13). Controls were then analyzed to ensure only background fluorescence for WT dystroglycan without peptide tags and a detectable signal only for the proper tag for the Flag-Dystroglycan and HA-Dystroglycan only constructions. All other samples were then exported and the ratio of APC-FITC (proteolysis marker) was divided by the FITC fluorescence (expression marker). The median ratio for the WT dystroglycan tagged with both Flag and HA was calculated. To calculate the fold change, each cell ratio was divided by the median ratio of the WT. Median fold change for each sample was then graphed using GraphPad Prism Software Version 9.4.1 where each dot represents a biological replicate.

### RAD TGT Assay

Force measurements were carried out using rupture and deliver tension gauged tether assay as previously described (68). Briefly, cells were plated in a 24 well plate and, once between 70-90% confluent, transfected as above. On the day of transfection, a 96 well glass bottom plate was prepared by plating first biotin-BSA (Thermo) and fibronectin (Sigma), followed by neutravidin (Thermo), followed by annealed 12 pN or 54 pN fluorescent TGTs (IDT), bound to WDV-Echistatin. The plate was incubated at 4°C in the dark overnight and the cells were incubated at 37°C with 5% CO2 overnight. The day after transfection, cells were unadhered from the plate, washed, and resuspended in OptiMem. They were counted and 15,000 cells were plated into two wells, one with 12 pN TGTs and the other containing 54 pN TGTs. Cells were incubated for 2-4 hours at 37°C in the dark, trypsinized, and resuspended in flow buffer (1X DPBS (Corning) supplemented with 1% BSA and 1 mM EDTA). Samples were kept on ice in the dark until run on BD Accuri C6 Plus Personal Flow Cytometer using BD Accuri C6 Plus (64-bit) Software Version 1.034.1. Analysis was done using FlowJo Version 10.8.1 and samples were gated for single cells with a APC fluorescent signal (Figure S14). For analysis on only GFP+ cells, samples were also gated for positive FITC fluorescence relative to nontransfected cells.

To graph samples, fold change compared to the WT-GFP sample was calculated by dividing all cells APC fluorescence (top strand of the TGT) by the median APC fluorescence of WT-GFP. In this way WT-GFP was always a fold change of 1 and any value above has higher signal which correlates to higher force and any value below 1 has a low signal and lower force. Median fold change was graphed using GraphPadPrism Version 9.4.1 where each point represents a biological replicate.

### Wound Healing

NIH-3T3 cells were reverse transfected with dystroglycan constructs or empty vector and plated in duplicate in a fibronectin coated, 24 well plate. After 36 hours, a scratch was made using a P200 pipette tip, the cells were washed once with DPBS, and media was replaced with DMEM supplemented with 5% FBS. Images were taken at respective time points using an EVOS FL Auto microscope. Four image areas were marked and monitored for each well, and wound are measurements were performed independently with Adobe Photoshop and the MRI Plugin for ImageJ. The data from four independent experiments were analyzed and plotted using GraphPad Prism. Each wound area was normalized where time=0 represented 100 percent of the area. The normalized data was then used for a two-way ANOVA analysis to determine statistical significance.

## SUPPLEMENTAL METHODS

### Analytical Ultracentrifugation

Dystroglycan protein was purified as above, except that after size exclusion chromatography, it was concentrated to an absorbance of 0.6 at 280 nm. Sedimentation velocity AUC experiments were performed in an Optima Analytical Ultracentrifuge (Beckman) with 2-channel Ti centerpieces (Nanolytics), quartz windows (Beckman), and An-60 rotor (Beckman). Cells were centrifuged at 50k rpm at 20°C for >9 hours with absorbance measured at 280 nm. Data analysis was completed using UltraScan III Software Version 4.0 (95) and the protocol published by the software developers (based on this method (96)). AUC experiments were done with two biological replicates which were run in duplicate to ensure accuracy.

### Evolution Conservation Analysis

Analysis of dystroglycan evolution conservation and creation of the logo plots were all done through a Python script available on the Gordon lab GitHub (https://github.com/AdamTSmiley). Dystroglycan amino acid sequences from multiple species were downloaded from UniProt using the accession codes listed in Table S2. They were aligned using Clustal Omega through the European Bioinformatics Institute (97). A simple conservation score was then calculated by determining the frequency of the most common amino acid in that position. Logo plots were created by calculating the frequency of each amino acid for each position.

## ACKNOWLEDGEMENTS

We would like to thank Jim Ervasti for helpful discussions. This study was supported by an NIH NIGMS R35 GM119483 grant to WRG and R35 GM118047 to HA. MJMA was supported by NIH T32 GM008244 (UMN MSTP) and T32 AR007512 (UMN Muscle Training). ANH received salary support from AHA Fellowship 17PRE33330000. ATS received salary support from NIH T32 GM132029 (Chemistry-Biology Interface Training Grant). We’d like to thank the staff at the University of Minnesota University Imaging Centers (SCR_020997) and Minnesota Nano Center (Award Number ECCS-2025124) for the use of their facilities and their helpful discussions. Thank you especially to Matthew Johnson at the Minnesota Nano Center for his help with AUC. Use of the Advanced Photon Source was supported by the U.S. Department of Energy under contract No. DE-AC02-06CH11357 and the Northeastern Collaborative Access Team, which is supported by NIH NIGMS P30 GM124165. Some images were created using BioRender.com. WRG is a Pew Biomedical Scholar.

## AUTHOR CONTRIBUTIONS

MJMA, ANH, ATS, MRP, and WRG designed experiments. MJMA, ANH, and KS acquired data. MJMA, KS, EG, LG, RLE and CU processed and refined the X-ray crystallography structure. MJMA, ANH, ATS, MRP, EJA, RLE, CU, and WRG analyzed and interpreted data. MJMA, ANH, EJA, and WRG wrote the manuscript. All authors edited the manuscript.

